# Orthosteric and allosteric effects of anti-CRISPR II-C1 inhibition on *Geo*Cas9 from integrated structural biophysics

**DOI:** 10.64898/2026.04.08.717222

**Authors:** Alexa L. Knight, Helen B. Belato, Charlotte S. Dresser, Chinmai Pindi, Briana J. Mercado, Praise Lasekan, Jinping Luo, Pablo R. Arantes, Gerwald Jogl, Giulia Palermo, George P. Lisi

**Affiliations:** Department of Molecular Biology, Cell Biology & Biochemistry, Brown University, Providence, RI USA; Departments of Bioengineering & Chemistry, University of California Riverside, Riverside, CA USA; Transgenic & Genome Editing Facility, Brown University, Providence, RI USA; Giuliani RNA Center, Brown University, Providence, RI USA

**Author notes:** Co-first authors (Alexa L. Knight, Helen B. Belato). Present address: Department of Chemistry, Brown University, Providence, RI USA.

## Abstract

Anti-CRISPRs (Acrs) are small protein inhibitors of CRISPR-Cas effectors that originate from the translated genetic material of bacteriophage. Harnessing the natural ability of Acrs to bind and disrupt CRISPR-Cas editing can provide enhanced spatiotemporal control of gene editing. Recent studies have revealed diverse structures and functions of Acrs, however, atomistic studies of the specific molecular mechanisms behind Acr inhibition are lacking. Here, we reveal how structure, function, and dynamics govern AcrIIC1 inhibition of Cas9 from *G. stearothermophilus* (*Geo*Cas9) via its HNH nuclease domain. An X-ray crystal structure of the *Geo*HNH-AcrIIC1complex reveals a conserved binding interface at the catalytic site and disruption of crucial electrostatic contacts known to modulate the thermostability of *Geo*Cas9. AcrIIC1 binding also rewires the intrinsic dynamics of the *Geo*HNH domain, stimulates millisecond motions that are absent from the unliganded nuclease, and attenuates the guide RNA affinity of *Geo*Cas9. Subsequent AcrIIC1 mutations in residues at its crystallographic binding interface uncouple Acr binding from inhibition, providing new insight into mechanism by which AcrIIC1 acts on *Geo*Cas9.

## Introduction

Applications of CRISPR-associated protein 9 (CRISPR-Cas9) have expanded from a biological tool for exploring gene editing to a promising therapeutic for human disease, including recent efforts to mitigate antibiotic resistance in bacterial infections(1–6). However, spatiotemporal control over CRISPR-Cas function remains a primary limitation, leading to off-target effects that could lead to large-scale genomic instability(7, 8). Specificity-enhancing mutations and directed evolution pipelines have mitigated this issue, but many of these insights have yet to translate to Cas effectors that function outside of mesophilic regimes(9–11). An emergent thermophilic Cas9 from *Geobacillus stearothermophilus* (*Geo*Cas9) has an expanded temperature range for double stranded DNA cleavage of ≥75[°C as well as increased thermostability and longevity in human serum(12, 13). Recent work has characterized some aspects of the intricate mechanism of *Geo*Cas9, including that structural fluctuations on multiple timescales govern the allosteric communication between its spatially separated domains(12, 14, 15). Most recently, cryo-EM structures have revealed a critical role for the “wedge” (WED), between the RuvC nuclease and PAM-interacting domains, in DNA unwinding and mutations in WED have produced the first high-specificity *Geo*Cas9, termed i*Geo*Cas9(15, 16).

In addition to protein engineering campaigns to improve the specificity and efficiency of Cas effectors, exogenous means of spatiotemporal control have leveraged bacteriophage-encoded anti-CRISPR proteins (Acrs). Several groups have investigated the structure and biochemistry of Acrs, which emerged naturally from an “evolutionary arms race” to bind and inhibit Cas effectors by diverse mechanisms(17–22). Here we provide atomic details of an Acr originating from *Neisseria meningitidis*, AcrIIC1, and its complex with *Geo*Cas9. Prior work reported AcrIIC1 to be a broad-spectrum inhibitor of related Type II-C Cas effectors from *N. meningitidis* (*Nme*Cas9), *Campylobacter jejuni* (*Cje*Cas9), and *Geo*Cas9. While AcrIIC1 does exploit some of the conserved features between *Geo*Cas9 and *Nme*Cas9 for binding, we also report that AcrIIC1 inhibition of *Geo*Cas9 imparts a distinct modulation of its HNH nuclease dynamics and function(17). As a consequence, AcrIIC1 not only inhibits cleavage activity through steric occlusion and rewiring of *Geo*HNH flexibility, but also by tampering with the guide RNA affinity of the full-length *Geo*Cas9.

To visualize the structural and biochemical signatures of AcrIIC1 inhibition of *Geo*Cas9, we combined nuclear magnetic resonance (NMR), X-ray crystallography, and all-atom molecular dynamics (MD) simulations with biophysical and functional assays. We resolve the interface between *Geo*HNH and AcrIIC1 by X-ray crystallography, identifying residues S78 and C79 of AcrIIC1 that directly interact with D581 and H582 of *Ge*oHNH, respectively. We describe how AcrIIC1 binding to the HNH domain of *Geo*Cas9 modulates its structure and rewires its intrinsic motions. We demonstrate that beyond a direct interaction that sequesters key residues from the *Geo*HNH active site (orthosteric), AcrIIC1 propagates long-range structural perturbations through *Geo*HNH, modulates guide RNA affinity of *Geo*Cas9, and potentially binds a secondary (allosteric) location within the *Geo*HNH domain. To assess the contribution of individual amino acids to the binding interface, we engineered S78A, C79P, and S78A/C79P mutations within AcrIIC1. The Cys-to-Pro substitution at residue 79 was designed to simultaneously disrupt multiple interactions of AcrIIC1 with the *Geo*HNH backbone observed crystallographically. NMR spectra of ^15^N-*Geo*HNH with S78A AcrIIC1 recapitulate the perturbations of wild-type (WT) AcrIIC1, while C79P has no effect. NMR relaxation experiments point to a dynamic rewiring of *Geo*HNH, the extent of which is tuned by the AcrIIC1 mutations. Simiarly, functional studies show none of our mutant variants inhibit GeoCas9 cleavage activity. All-atom MD simulations of WT and mutant *Geo*HNH-AcrIIC1 complexes show that WT AcrIIC1 stabilizes the HNH catalytic loop, while AcrIIC1 mutants increase backbone flexibility and conformational heterogeneity. Consistent with functional studies, MD confirms that AcrIIC1 inhibition of *Geo*Cas9 requires both active-site occupancy and disruption of a coordinated dynamic network within *Geo*HNH, with variants suggesting that subtle perturbations to the protein-protein contacts can partially uncouple binding from inhibition.

## Results

### A crystal structure of the *Geo*HNH-AcrIIC1 complex reveals a conserved binding interface

To elucidate the binding mode of AcrIIC1 to *Geo*Cas9, we determined a 1.7 Å resolution X-ray crystal structure of AcrIIC1 bound to the HNH nuclease of *Geo*Cas9. Aligning almost identically with a previously published crystal structure of AcrIIC1 bound to *Nme*HNH (**Figure S1**), the inhibitor directly interacts with the active site of *Geo*HNH. Crucially, the active site residues D581 and H582 form hydrogen bonds with AcrIIC1 residue S78, as well as the sidechain and backbone amide of C79 (**Figure 1A**). Second sphere charged residues of *Geo*HNH further stabilize the inhibitor. Structural studies of closely related HNH domains, including the *Nme*HNH that shares 44% sequence identity with *Geo*HNH, showed similar charge stabilization of Acrs via identical second sphere residues (**Figure 1B,C**)(17, 21).

**Figure 1.**
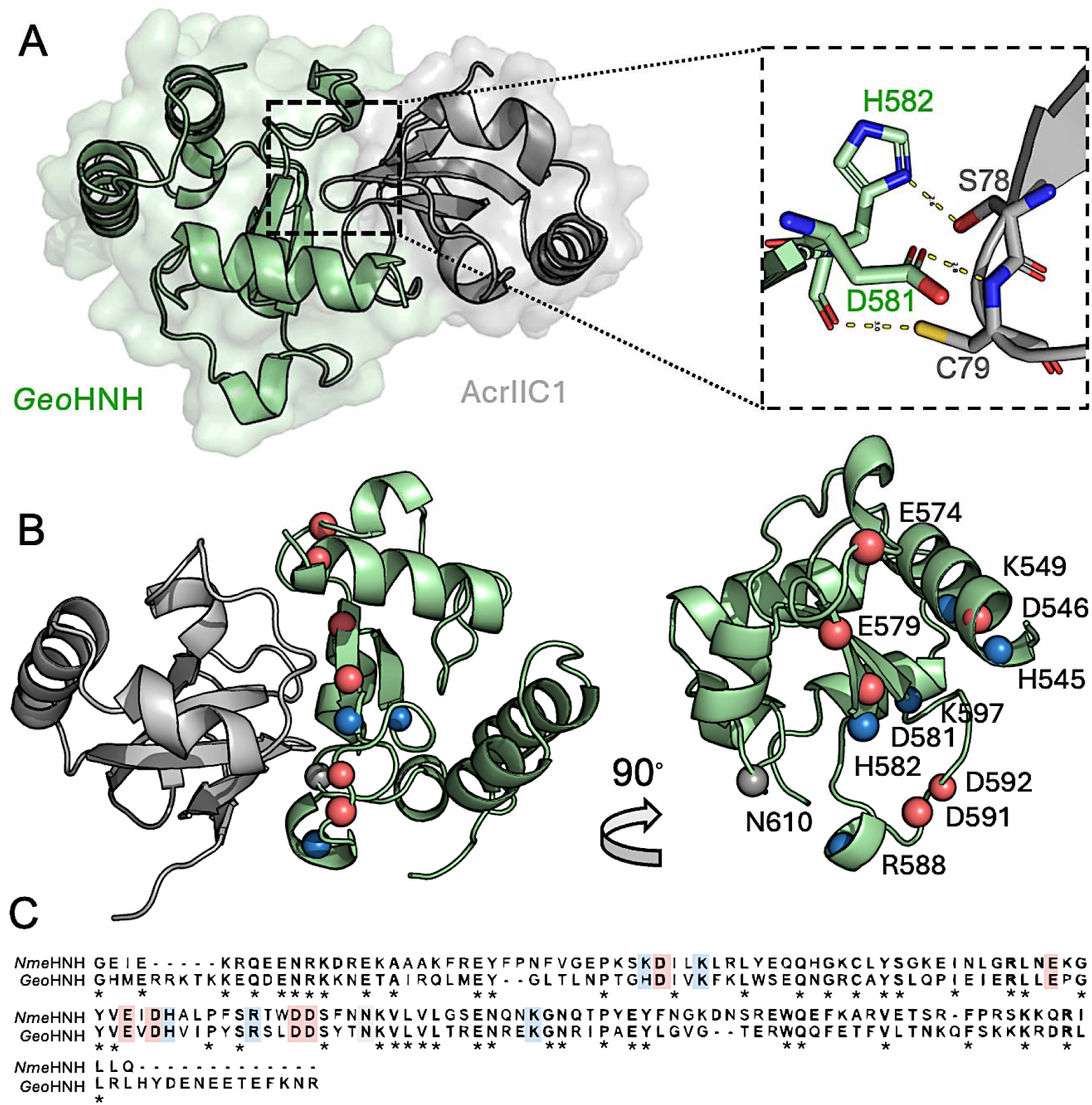
(**A**) X-ray crystal structure of the *Geo*HNH-AcrIIC1 complex. The inset highlights a direct interaction of AcrIIC1 with the *Geo*HNH catalytic residues. (**B**) The X-ray structure positions conserved charged residues at the HNH-Acr interface to stabilize the complex, as in the AcrIIC1 complex with *Nme*HNH, a *Geo*Cas9 ortholog. (**C**) Sequence alignments of *Nme*HNH and *Geo*HNH show a high degree of conservation at residues forming the protein-protein interface with AcrIIC1. Charged residues highlighted in (B) are colored and other conserved residues are denoted by asterisks.

### Solution NMR studies of the *Geo*HNH-AcrIIC1 complex reveal direct and long-range structural perturbations

To assess the effect of AcrIIC1 on the atomic structure of *Geo*HNH, we first used NMR to visualize the *Geo*HNH amide backbone structure in response to increasing concentrations of AcrIIC1. ^1^H-^15^N transverse relaxation optimized spectroscopy-heteronuclear single quantum coherence (TROSY-HSQC) spectra of *Geo*HNH revealed that AcrIIC1 binding induced chemical shift perturbations (CSPs) throughout the nuclease domain, despite interacting with only two residues in the X-ray crystal structure. We observed a large majority of *Geo*HNH resonances to be in slow exchange between the free and bound states of *Geo*HNH when titrated with wild-type (WT) AcrIIC1, suggesting a tight binding interaction (**Figure 2A, Figure S2)**. The greatest magnitude and concentration of these CSPs are observed near the *Geo*HNH catalytic site, consistent with our crystal structure, though AcrIIC1-induced CSPs propagate well into the core of *Geo*HNH (**Figure 2B,C**). Importantly, a reciprocal NMR titration using ^15^N-labeled AcrIIC1 produced a similar degree of slow-exchange CSPs occurring predominantly at the expected crystallographic interface (**Figure S3)**.

**Figure 2.**
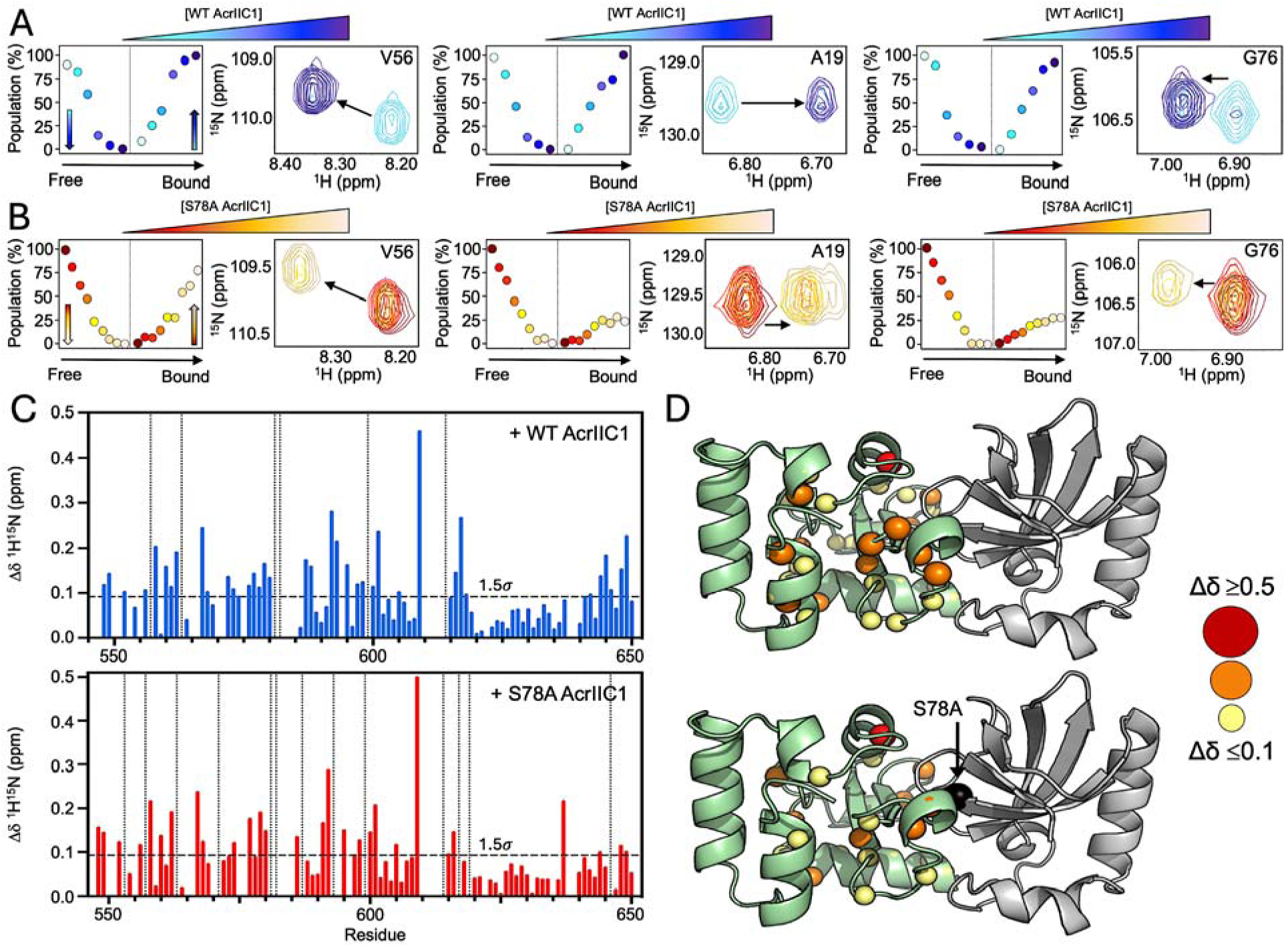
Snapshots of 1H-15N HSQC titrations of *Geo*HNH with AcrIIC1 inhibitor. *Geo*HNH is titrated with **(A)** WT AcrIIC1 (light-to-dark blue) and **(B)** S78A AcrIIC1 (dark-to-light red). Slow exchange CSPs suggest tight binding and populations of free and AcrIIC1-bound *Geo*HNH are derived from resonance volumes shown as dot plots. The inflection between ligation states is indicated by vertical dashed lines. **(C)** Per-residue NMR CSPs of *Geo*HNH caused by WT AcrIIC1 (upper, blue bars). The same concentration of S78A AcrIIC1 induces very weak CSPs in *Geo*HNH (lower, black bars), while further addition of S78A AcrIIC1 rescues the CSPs (red bars). Dashed vertical lines denote sites of NMR line broadening in each plot. **(D)** CSPs >1.5σ of the 10% trimmed mean of all shifts (horizontal dashed line in (B)) are mapped onto the *Geo*HNH structure as spheres, where the size and color correlate with intensity of the CSP. The S78A mutation is noted with a black

To validate the contribution of individual AcrIIC1 residues depicted in the X-ray structure to the *Geo*HNH binding interaction, we introduced single-point mutations to S78 (S78A) and C79 (C79P, to disrupt both the Cys side chain and amide backbone interactions) of AcrIIC1. Circular dichroism and ^1^H-^15^N NMR spectra confirm that S78A and C79P variants remain folded with intact secondary structure (**Figure S4)**. The S78A AcrIIC1 variant induced a WT-like effect on *Geo*HNH in ^1^H-^15^N TROSY-HSQC titrations (**Figure 2B-D**), though we noted additional line broadening with S78A AcrIIC1. The C79P AcrIIC1 variant induced no CSPs in *Geo*HNH, highlighting the hydrogen bonding between the sidechain and backbone of C79 with *Geo*HNH as crucial for maintaining the complex. Additionally, we engineered a S78A/C79P negative control variant and also observed no CSPs in the ^1^H-^15^N NMR spectrum of *Geo*HNH.

To further assess the interaction between *Geo*HNH and WT AcrIIC1 or variants, we measured *R*_1_, *R*_2_, and ^1^H-[^15^N] heteronuclear NOE relaxation parameters that are sensitive to fast timescale conformational fluctuations and report on global tumbling (**Figure S5**). When compared to *Geo*HNH alone, *Geo*HNH in complex with both WT and S78A AcrIIC1 showed globally depressed *R*_1_ and elevated *R*_2_ values consistent with an increase in molecular weight. As expected from initial NMR titrations, *R*_1_ and *R*_2_ values of *Geo*HNH in the presence of C79P AcrIIC1 showed no change from the apo protein. ^1^H-[^15^N] NOEs of *Geo*HNH in the presence of WT or S78A AcrIIC1 revealed some local fluctuation in backbone motion due to AcrIIC1, while the C79P NOE profile resembled that of apo *Geo*HNH. We then used these data to calculate per-residue order parameters (*S*^2^) for the free and Acr-bound states of *Geo*HNH (**Figure 3A, Figure S6)**. Most notably, regions in close sequence and spatial proximity to the *Geo*HNH active site experienced sharp increases in *S*^2^ in the presence of AcrIIC1, suggesting the inhibitor suppresses *ps-ns* timescale flexibility in these regions. Collectively, NMR titrations and relaxation experiments point to the interactions of AcrIIC1 residue C79 with *Geo*HNH as essential to the inhibited complex, while S78 plays a lesser role. Additionally, we observed a suppression of pico-nanosecond (*ps-ns*) timescale flexibility within *Geo*HNH, which has previously been shown to control nuclease function and allostery(12).

**Figure 3.**
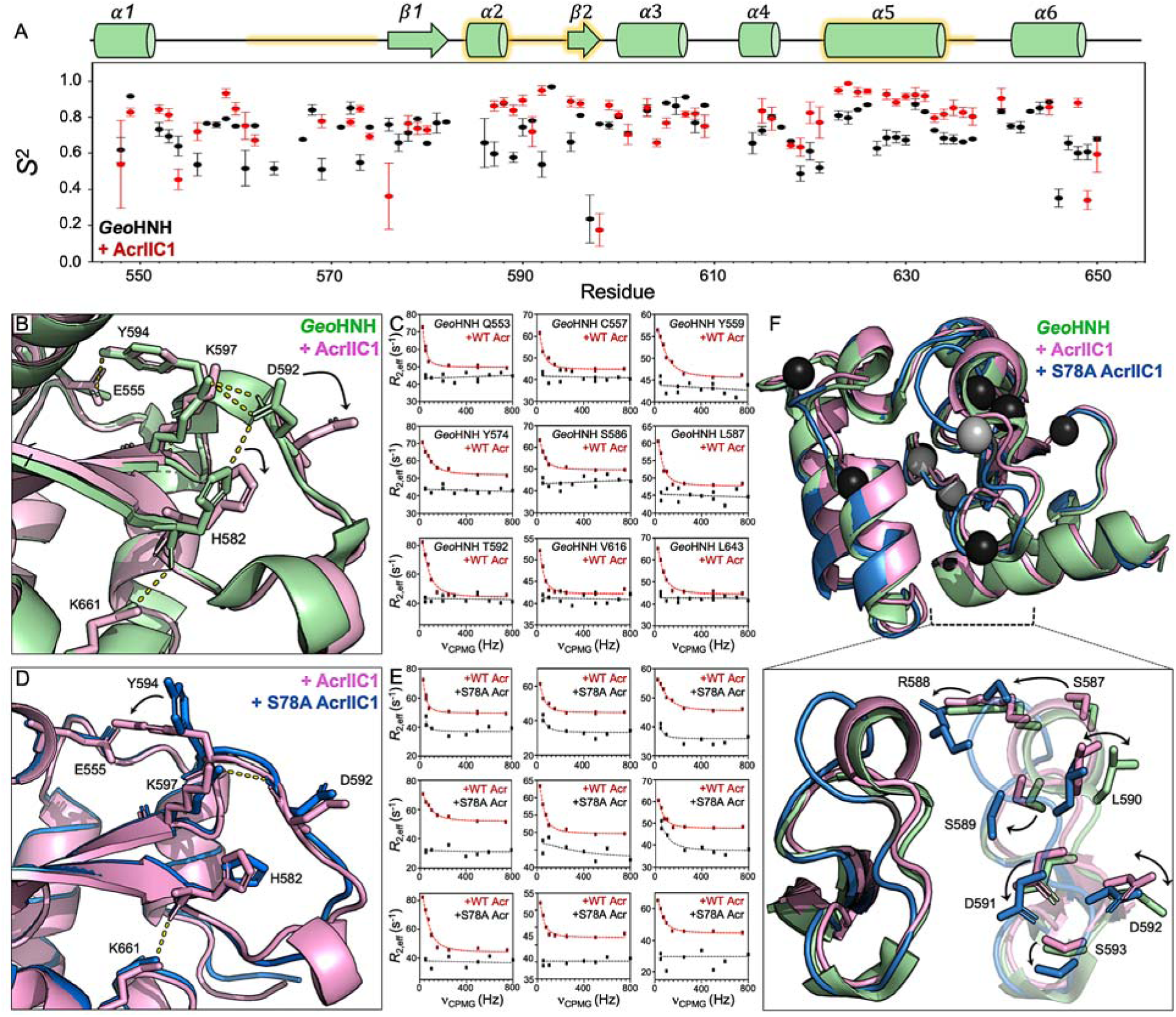
(**A**) NMR order parameters (S2) of *Geo*HNH in the absence (black) and presence (red) of AcrIIC1. A cartoon of *Geo*HNH structure (top) highlights AcrIIC1-induced suppression of fast timescale flexibility (yellow glow). **(B)** X-ray crystal structure of *Geo*HNH (green, PDB: 7MPZ) overlaid with that of AcrIIC1-bound *Geo*HNH (pink, PDB:10VC). AcrIIC1 disrupts electrostatic contacts within a salt bridge network (dashes) via displacement of key catalytic side chains (arrows). **(C)** CPMG relaxation dispersion profiles of *Geo*HNH (black, flat profiles) and AcrIIC1-bound *Geo*HNH (red curves), highlighting the stimulation of *ms* motions upon disruption of the salt bridge network. **(D)** X-ray crystal structures of WT (pink, PDB: 10VC) and S78A AcrIIC1-bound *Geo*HNH (blue, PDB: 10VB) are nearly identical at the *Geo*HNH active site. **(E)** CPMG relaxation dispersion experiments indicate that the S78A mutation limits the ability of AcrIIC1 to stimulate *ms* motions in *Geo*HNH. **(F)** Sites of *ms* timescale flexibility are mapped onto a structural overlay of *Geo*HNH (green, PDB: 7MPZ), AcrIIC1-bound *Geo*HNH (pink, PDB: 10VC), and S78A AcrIIC1-bound *Geo*HNH (blue, PDB: 10VB). Black spheres indicate flexible residues in the WT Acr-bound complex, while gray spheres are flexible residues in the S78A Acr-bound complex. CPMG relaxation dispersion localizes to a region of crystallographic structural heterogeneity among the three *Geo*HNH domains, highlighted for the backbone and side chains.

### AcrIIC1 disrupts salt bridge networks of *Geo*HNH, stimulating millisecond timescale motions

In a prior study of *Geo*HNH, we analyzed an active site salt bridge network and showed that mutation of a central residue (K597) disrupted many charged contacts, leading to significant structural and functional perturbations(12). Namely, a K597A mutation induced widespread and globally synchronous millisecond (*ms*) timescale motions that were absent from WT *Geo*HNH. Further, these motions rewired *Geo*HNH such that a single Lys-to-Ala mutation enhanced the thermostability of full-length *Geo*Cas9 and increased its functional range for DNA cleavage. Our X-ray crystal structure of the *Geo*HNH-AcrIIC1 complex reveals the same extent of salt bridge disruption. Stabilizing contacts between K597, H582, and D592 are now broken, allowing the loop containing the catalytic histidine to flip outward, while K597 rotates from the *Geo*HNH surface toward the core of the protein (**Figure 3B)**. We then interrogated the impact of these disruptions on *Geo*HNH flexibility with Carr-Purcell-Meiboom-Gill (CPMG) relaxation dispersion NMR experiments, noting that *ms* motions, as in our prior work, were now apparent in *Geo*HNH (**Figure 3C**). Ten residues in the *Geo*HNH-AcrIIC1 complex now display curved relaxation dispersion profiles, while no such profiles exist in *Geo*HNH alone. Residues undergoing chemical exchange on this timescale are not isolated to the binding interface with AcrIIC1, but rather span the length of the *Geo*HNH, including sites of interaction with nucleic acids and the recognition (REC) domain within *Geo*Cas9. This suggests the potential for AcrIIC1 to alter the communication between REC and HNH, which has been shown to be crucial for coordinating double stranded DNA cleavage(14, 23–25). An assessment of the S78A AcrIIC1 variant showed a nearly identical active site arrangement of the inhibitor bound state, though the extent of *ms* motion was not as pronounced (**Figure 3D,E**). Interestingly, the region of *Geo*HNH harboring the bulk of its *ms* motion and greatest decreases in *ps-ns* motion, also displays conformational heterogeneity between our *Geo*HNH, WT AcrIIC1-bound, and S78A AcrIIC1-bound crystal structures (**Figure 3F**) which may explain the subtle differences in NMR-detected flexibility.

### Mutations at the crystallographic binding interface limit the ability of AcrIIC1 to inhibit the guide RNA interactions and DNA cleavage of *Geo*Cas9

We next tested the functional consequences of mutations at the crystallographic *Geo*HNH-AcrIIC1 binding interface with *in vitro* DNA cleavage assays of *Geo*Cas9 in the presence of WT, S78A, C79P, or S78A/C79P AcrIIC1. WT AcrIIC1 inhibits target DNA cleavage by *Geo*Cas9 in a concentration dependent manner, consistent with prior reports that also demonstrated a significant molar excess of AcrIIC1 being required for inhibition (17). At similar concentrations, we observed essentially no inhibition by the S78A, C79P, or S78A/C79P AcrIIC1 variants (**Figure 4A,B**). We also noted that the order in which components were added to the reaction mixture was critical to the efficiency of inhibition. When the *Geo*Cas9 ribonucleoprotein complex (RNP) was formed prior to the addition of AcrIIC1, little-to-no inhibition was observed (**Figure S7)**. This suggests that sgRNA-induced conformational changes, which typically suppress the flexibility of Cas enzymes, may occlude the AcrIIC1 binding site.

**Figure 4.**
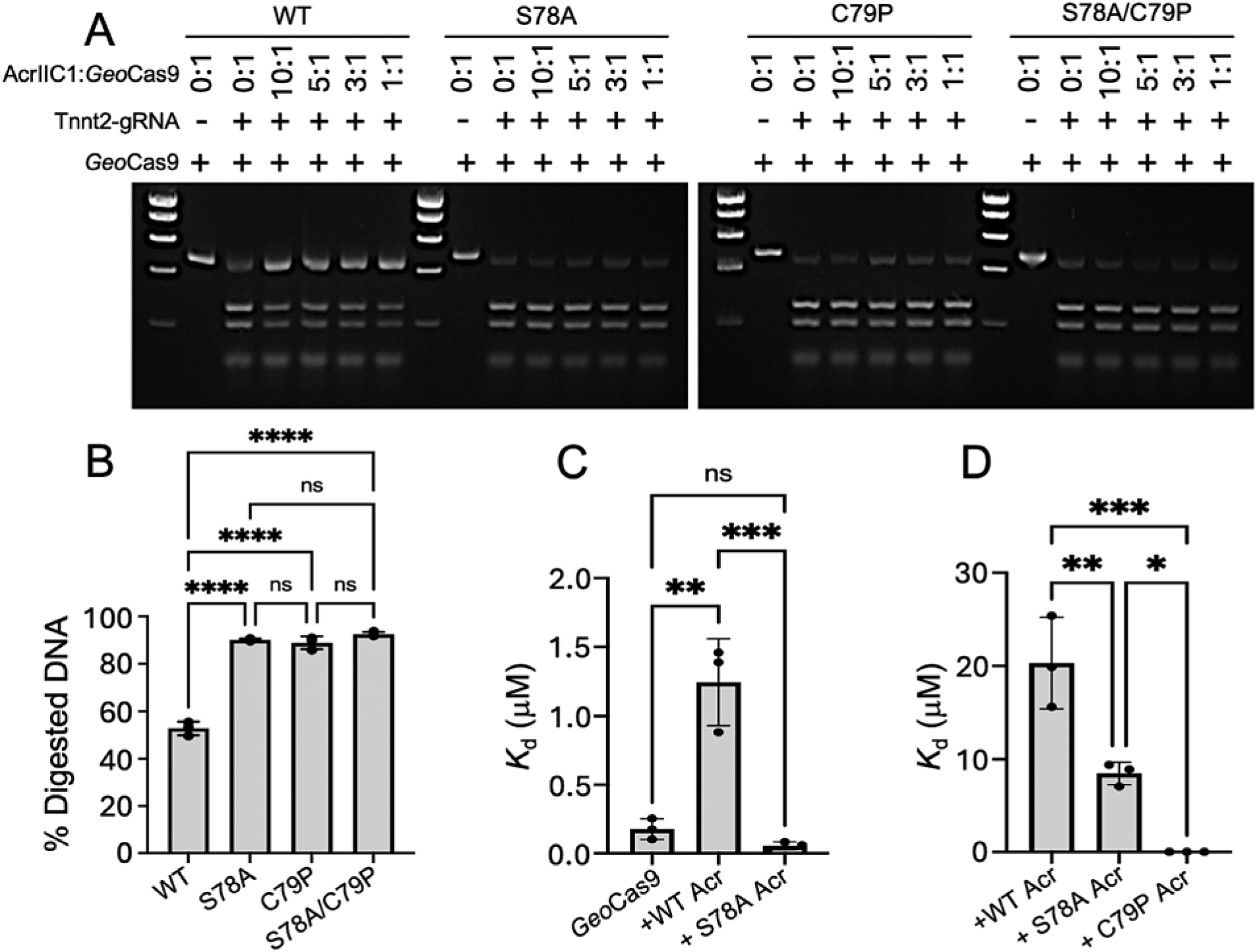
(**A**) *In vitro* DNA cleavage assays indicate that AcrIIC1 mutations at the crystallographic binding interface ablate its inhibition. *Geo*Cas9 was incubated with varying concentrations of WT, S78A, C79P, and S78A/C79P AcrIIC1, after which RNPs were formed by incubating the AcrIIC1-bound *Geo*Cas9 with sgRNA and assayed for target DNA cleavage at 37 °C. **(B)** Bar graph quantifying the DNA cleavage via band intensities from the agarose gel. ****p<0.0001. n=3 technical replicates for each sample. **(C)** MST-derived *K*d values of *Geo*Cas9 for its sgRNA in the absence and presence of WT and S78A AcrIIC1. **p<0.0011; ***p<0.0006. n=3 technical replicates for each sample. **(D)** MST-derived *K*d values of *Geo*HNH for WT, S78A, and C79P AcrIIC1. *p<0.0282; **p<0.0061; ***p<0.0004. n=3 technical replicates for each sample.

These data led us to wonder if the affinity of *Geo*Cas9 for its single-guide (sg) RNA was modulated by the inhibitor. We used microscale thermophoresis (MST) to measure binding affinities between *Geo*Cas9 and a Cy5-labeled sgRNA from a recently published cryo-EM structure (PDB:8UZA), using *Geo*Cas9 alone or in the presence of WT or S78A AcrIIC1. Consistent with functional assays, MST showed that WT AcrIIC1 reduced the affinity of *Geo*Cas9 for its sgRNA (*K*_d_ = 124 ± 91 nM for *Geo*Cas9 compared to *K*_d_ = 1.2 ± 0.4 μM for *Geo*Cas9-AcrIIC1, **Figure 4C**). Surprisingly, we found that despite WT-like X-ray structures and CSPs in NMR titrations of *Geo*HNH with S78A AcrIIC1, this variant did not attenuate the affinity of *Geo*Cas9 for sgRNA (*K*_d_ = 52.2 ± 23.7 nM). These data are also consistent with functional assays, which show little-to-no inhibition from the S78A AcrIIC1 variant. Interestingly, we found that MST assessing protein-protein interaction between *Geo*HNH and WT or S78A AcrIIC1 showed little difference in binding affinity between the proteins themselves **(Figure 4D),** with S78A returning a slightly tighter *K*_d_ than WT AcrIIC1 to *Geo*HNH (*K*_d_ = 19.9 ± 12.5 μM and 7.93 ± 4.5 μM, respectively). Despite its structural impact in NMR titrations, differences between an isolated *Geo*HNH domain and full-length *Geo*Cas9 suggest that direct side chain-specific interactions between S78 of AcrIIC1 and the catalytic H582 is in fact required for inhibition.

### MD simulations pinpoint rewired *Geo*HNH-AcrIIC1 contacts that destabilize the binding interface

To complement our experimental structural studies, we performed extensive all-atom molecular dynamics (MD) simulations of the *Geo*HNH-AcrIIC1 complex to investigate how AcrIIC1 (and the interfacial variants) reshaped the conformational landscape of *Geo*HNH. Simulations were carried out for the *Geo*HNH-AcrIIC1 complex as well as the S78A, C79P, and S78A/C79P variants in triplicate, enabling robust quantitative comparisons of structural stability, flexibility, and allosteric coupling.

Across all replicates, the *Geo*HNH-WT AcrIIC1 complex remained structurally stable, reflected by low backbone root-mean-square deviations (RMSDs) relative to the crystallographic structure (**Figure S8A**). The variants exhibited RMSDs values comparable to WT, with S78A and C79P exhibiting slightly increased variability. Per-residue root-mean-square fluctuation (RMSF) analysis revealed higher intrinsic flexibility of *Geo*HNH than AcrIIC1 (**Figure 5A**). While the S78A variant largely preserved WT-like backbone dynamics, both the C79P and the S78A/C79P double mutant complexes exhibited elevated fluctuations (**Figure 5A**). Notably, increased fluctuations extended beyond the immediate binding interface, consistent with partial propagation of local perturbations into broader regions of *Geo*HNH, consistent with NMR CSP analysis.

**Figure 5.**
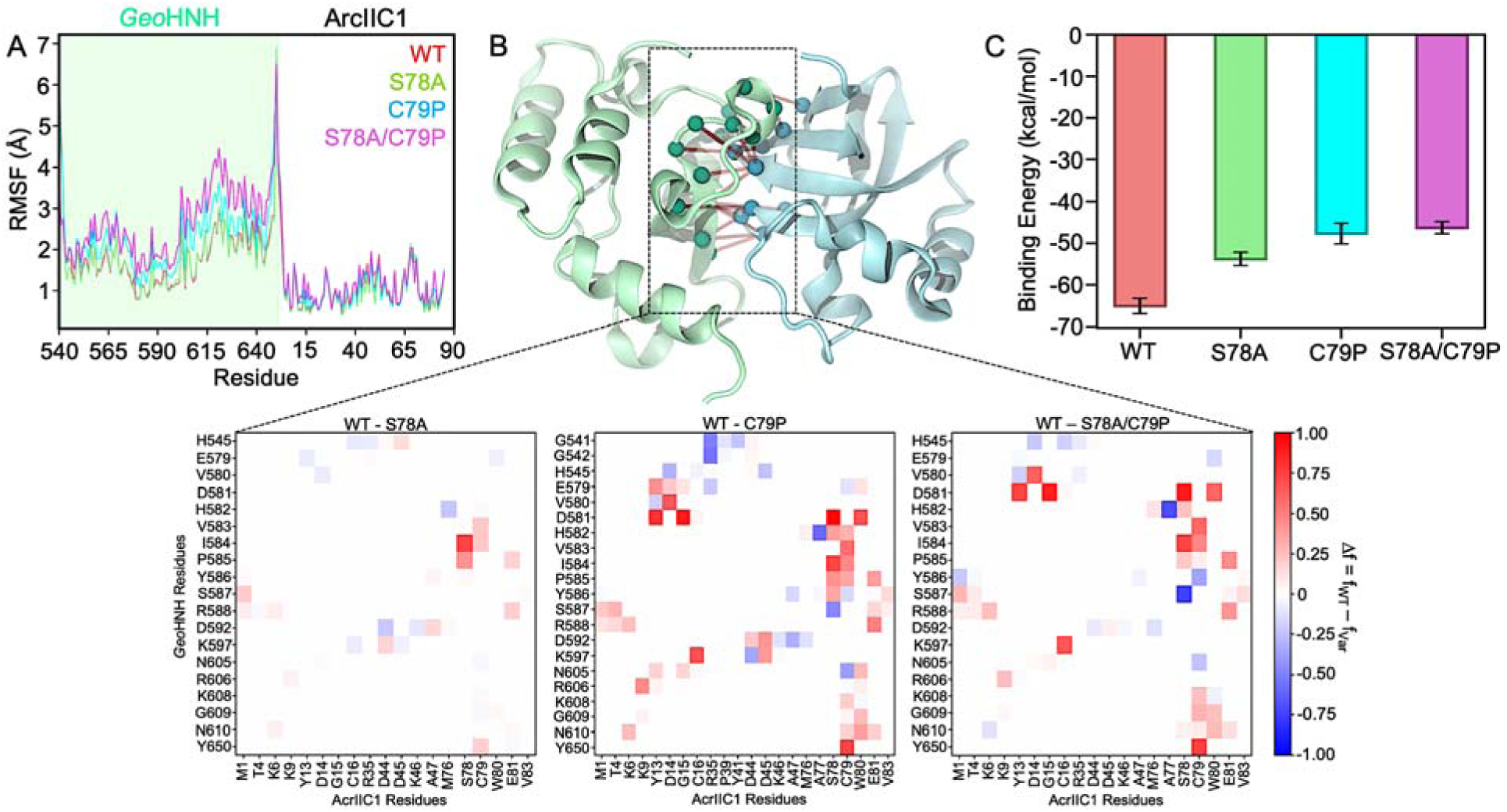
Effects of AcrIIC1 mutations on structural stability, binding energetics, and interface integrity of the *Geo*HNH-AcrIIC1 complex. **(A)** Per-residue root-mean-square fluctuations (RMSF) of the *Geo*HNH-AcrIIC1 complex across WT and mutant systems (S78A, C79P, and S78A/C79P). The shaded green region highlights the *Geo*HNH domain within the complex. **(B)** Representative structural snapshot of the *Geo*HNH-AcrIIC1 complex highlighting the primary binding interface. Spheres denote key residues participating in stable interfacial interactions (red lines) in the WT complex, corresponding to contacts enriched in WT (red squares) identified from the differential contact maps. Lower panels depict residue-residue contact difference maps (Δf) for S78A, C79P, and S78A/C79P variants relative to WT (Δf = WT − variant). Positive values (red) indicate contacts enriched or stable in the WT complex, whereas negative values (blue) indicate contacts newly formed in the mutant complexes. **(C)** Binding energies (kcal/mol) between *Geo*HNH and WT or mutant AcrIIC1 systems, showing reduced binding energy upon mutation (± standard error of mean).

To further assess the stability of the *Geo*HNH-AcrIIC1 complex, we calculated the total number of contacts formed between the proteins. Here, a contact is defined using a distance cut-off of 4.5 Å between heavy atoms of *Geo*HNH residues and AcrIIC1 residues. The WT complex maintained the highest number of intermolecular contacts, whereas S78A showed a modest reduction, and both C79P and S78A/C79P exhibited a substantial loss of contacts (**Figure S8B**). These observations are consistent with subtle disruptions to interfacial packing rather than complete dissociation of the complex. We also generated differential residue-residue contact maps, charting intermolecular interactions relative to the WT complex (**Figure 5B, S8B**). The WT complex is characterized by a dense network of persistent contacts at the *Geo*HNH-AcrIIC1 interface, several of which are substantially weakened or lost in the variant systems. These disrupted interactions are reflected as regions of positive Δf, highlighting key interaction hotspots that are sensitive to mutation. While some compensatory contacts emerge upon mutation of the primary interface (negative Δf), these are sparse and do not recapitulate the cohesive interaction network observed in the WT complex. Mapping of the WT-enriched contacts onto the complex structure reveals that residues surrounding S78 and C79 form a central interaction hub, underscoring their importance in maintaining interface integrity (**Figure 5B, S8B**).

Consistent with these structural and dynamical changes, interaction energy calculations revealed a progressive reduction in binding energy upon mutation. The WT complex exhibited the most favorable interaction energy, followed by S78A, while C79P and the S78A/C79P double mutant showed significantly weaker binding (**Figure 5C**). Together, these results demonstrate that while S78A induces only modest perturbations, the C79P and S78A/C79P variants substantially disrupt interfacial packing, increase conformational flexibility, and weaken binding, thereby compromising the inhibitory efficiency of AcrIIC1 against *Geo*HNH.

Collectively, the MD simulations indicate that the conformational heterogeneity of the *Geo*HNH-AcrIIC1 interface is heightened in the presence of mutations, which also destabilize the catalytic-loop geometry observed in the crystallographic complex. In agreement with X-ray and NMR analyses, effective inhibition by AcrIIC1 depends not only on active-site engagement but also on stabilization of coordinated dynamic interactions with *Geo*HNH. The behavior of the variants further suggests that subtle perturbations at the interface can partially decouple Acr binding from inhibition, highlighting the importance of dynamic allosteric regulation in this system.

These observations are consistent with functional assays, which reveal that the S78A AcrIIC1 variant exhibits little-to-no inhibitory activity despite retaining binding to *Geo*HNH. Notably, MST measurements of the *Geo*HNH-AcrIIC1 interaction show comparable binding affinities for WT and S78A AcrIIC1, with S78A having a slightly tighter apparent dissociation constant relative to WT (**Figure 4D**). This apparent discrepancy between binding affinity and inhibitory function suggests that the S78A mutation does not significantly impair complex formation but instead perturbs the functional coupling between binding and inhibition. In the context of our MD simulations, these results support a model in which S78A preserves overall binding but subtly alters interfacial dynamics and contact organization, leading to a loss of inhibitory efficiency without substantial destabilization of the complex. Together, the experimental and computational data highlight that effective inhibition by AcrIIC1 depends not only on binding affinity but also on the precise structural and dynamic integrity of the interface required to stabilize the catalytically inactive conformation of *Geo*HNH.

### NMR and AlphaFold3 modeling suggest a second binding location for AcrIIC1 on *Geo*HNH

We have noted in several prior studies that a shift in the intrinsic dynamics of *Geo*HNH to slower timescales is detrimental to its ability to propagate long-range allosteric information (14, 26). Balancing this point is our prior speculation, supported by functional assays, that despite this motional shift, mutation-induced disruption of salt bridges could modify or even strengthen contacts between HNH and the adjacent RuvC or REC domains of full-length *Geo*Cas9 (12). The influence of the AcrIIC1 binding partner, as opposed to a mutation, could be more complicated. As has been observed in related Cas9-Acr complexes, when our X-ray structure of *Geo*HNH-AcrIIC1 is overlaid with structures of sgRNA-bound or sgRNA:DNA-bound *Geo*Cas9, the steric clash is very high between AcrIIC1 and the RuvC domains in these static snapshots (**Figure S7**). However, slower timescale and larger amplitude motions of the *Geo*HNH domain could serve to accommodate the inhibitor at its primary binding site. Thus, AcrIIC1 may prefer to bind apo *Geo*Cas9 and prevent additional conformational dynamics that activate the nucleases.

AcrIIC1 caused large and widespread CSPs in NMR spectra of ^15^N-*Geo*HNH (**Figure 1A, S3**), which ceased at equimolar equivalents of AcrIIC1:*Geo*HNH. However, further titration to significant molar excesses of AcrIIC1 induced linear shifts in fast exchange at a small number of resonances **(Figure 6A)**. Distinct from the initial addition of AcrIIC1, these shifts map to two discrete regions of the *Geo*HNH domain **(Figure 6B)** that would be solvent exposed in the context of full-length *Geo*Cas9. We therefore speculated that *Geo*HNH may accommodate a second AcrIIC1 inhibitor at another interface. An AlphaFold3 model predicted a secondary binding location that aligned with one of the regions of NMR CSPs, with specific interactions between AcrIIC1 and the N605 and R606 sidechains, as well as the G618 backbone (27). Further, our AlphaFold3 model strongly resembles existing structures (PDB: 6J9N, 8IF0) of *Nme*HNH bound to both AcrIIC1 (identical to our crystallographic interface) and AcrIIC3, though the secondary Acrs have subtle differences in orientation (22, 28). NMR demonstrates the secondary interaction to be weak and/or transient, with an apparent *K*_d_ = 700 μM from a global fit of CSPs in fast exchange (**Figure 6C**). ^1^H-^13^C HMQC NMR experiments visualizing the histidine side chains of *Geo*HNH showed that in addition to expected line broadening of the catalytic H582 due to its direct orthosteric interaction with WT AcrIIC1, CSPs and/or line broadening of H551 and H659, which are distal to primary binding site, are also apparent (**Figure 6D**). H659 is especially close to the AlphaFold3-predicted secondary interface. It should be noted, however, that many CSPs from our NMR experiments also localized to the base of two α-helices on the face opposite the *Geo*HNH catalytic site (**Figure 6B**), making it a plausible surface for interaction.

**Figure 6.**
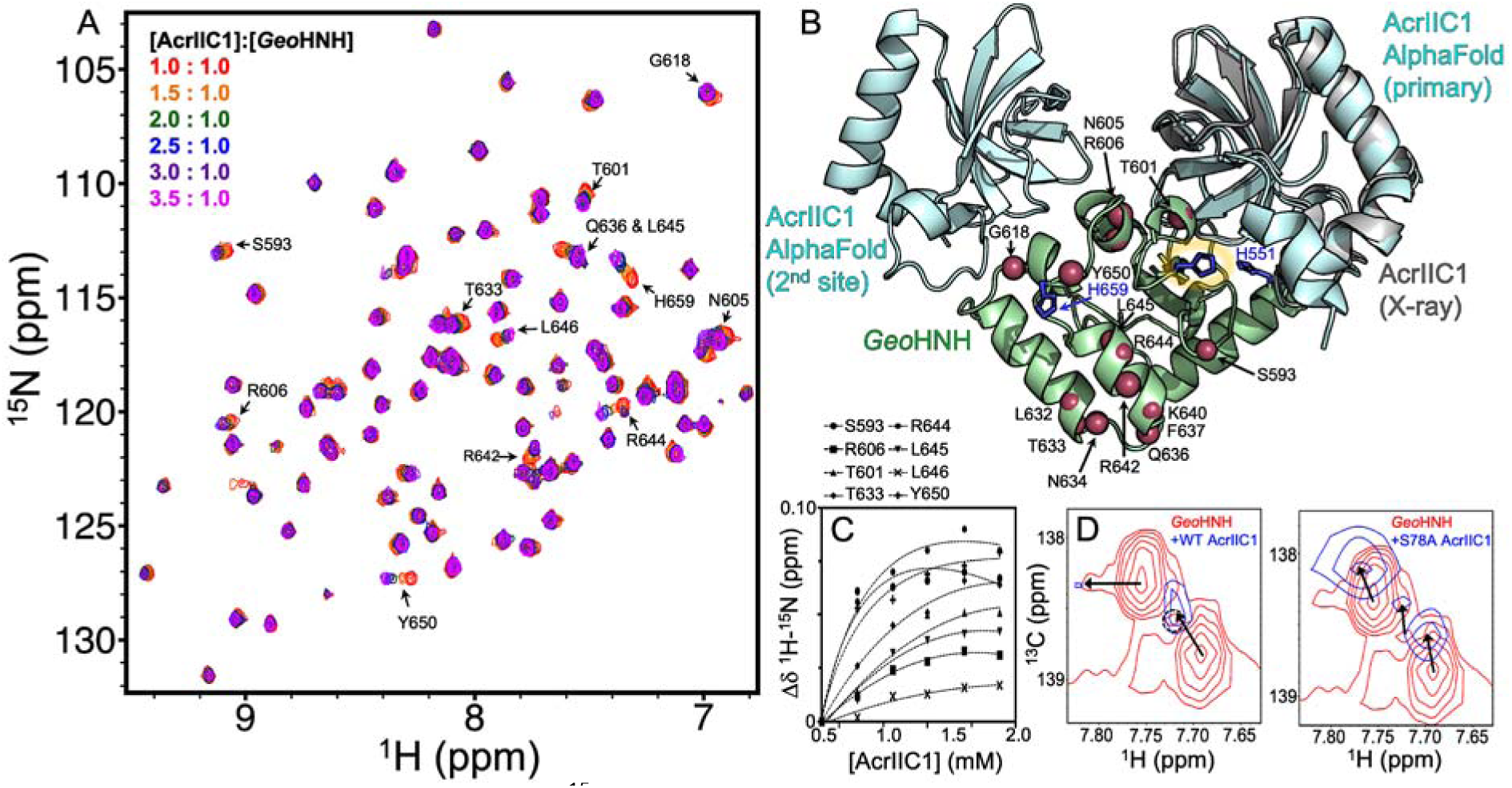
(**A**) NMR spectral overlay of 15N-*Geo*HNH revealing a small number of CSPs at super-saturating concentrations of AcrIIC1. The spectra are colored according to the Legend (inset). **(B)** AlphaFold3 model of *Geo*HNH (green) bound to two molecules of AcrIIC1 (cyan). The primary site matches our crystallographic interface (gray cartoon), while the second site localizes to regions of NMR CSPs (red spheres). Histidine side chains are shown as blue sticks and the catalytic H582 is circled. **(C)** Apparent binding affinity for the second site, based on a global fit of NMR chemical shift trajectories, is ∼ 710 μM. **(D)** 1H-13C NMR spectra of the His ε1 side chain show AcrIIC1-induced CSPs (arrows) or line broadening (circles) in all three native His residues of *Geo*HNH, despite two of these being outside the primary binding site.

## Discussion

Antil7CRISPRs have emerged as natural protein regulators of CRISPR-Cas effectors, but most mechanistic work has focused on mesophilic Cas systems and/or static views of Acr inhibition. Furthermore, previous biophysical studies of Acrs have generally focused on the structure of the Acrs themselves, rather than atomistic interactions with Cas proteins. Structural insights into several Cas9s in complex with AcrIIC1 have framed it as a broad-spectrum inhibitor sterically occluding a conserved HNH active site (17). Building on these prior insights, new atomic level information from NMR and MD simulations have refined a view of AcrIIC1 binding that reshapes the internal communication pathways within the thermophilic *Geo*Cas9 enzyme. This study presents evidence for a dynamic aspect of anti-CRISPR inhibition in which Acrs modulate the same networks within *Geo*HNH that govern RuvC and REC coordination and sgRNA affinity(26). Our prior work revealed that the core dynamics of *Geo*HNH are consistent up to temperatures of 40 °C, highlighting an evolutionary pressure to preserve fast motions under high entropic burden. We found no specific motional pathway comprised by *Geo*HNH residues, but rather a robust preservation of the internal dynamics themselves (26). We previously speculated that rewiring these core motions toward the slower *ms* timescale would increase the activity of *Geo*Cas9, as similar flexibility was found to be widespread and principal in the canonical *Sp*Cas9. However, such a rewiring of the *Geo*HNH domain by AcrIIC1 in fact has an inhibitory effect, highlighting a key distinction between the intrinsic dynamics that govern mesophilic and thermophilic Cas9, and thus the need to establish detailed, system-specific mechanistic rules for Cas effectors.

This study suggests that AcrIIC1 inhibits *Geo*Cas9 through a combination of orthosteric and allosteric effects that are more nuanced than pure activel7site steric hinderance. An X-ray crystal structure of the *Geo*HNH-AcrIIC1 complex highlights an interface that engages the HNH catalytic loop via AcrIIC1 residues S78 and C79, disrupting an electrostatic network within *Geo*HNH that rewires its local structure and dynamics(12). A region of conformational heterogeneity across several HNH crystal structures is observed, and NMR relaxation experiments reveal that AcrIIC1 suppresses *ps-ns* motions throughout this region (and others) while stimulating new, broadly distributed *ms* motion. Functionally, the interaction between *Geo*HNH residue H582 and S78 of AcrIIC1 is crucial to inhibition, while disruption of the D581-C79 interaction abolishes the binding of AcrIIC1 to *Geo*HNH and its inhibition of *Geo*Cas9. At high inhibitor concentrations, NMR and AlphaFold3 modeling indicate the existence of a weak secondary binding site on *Geo*HNH. We speculate that this secondary site could most plausibly affect closed conformations of sgRNA-bound *Geo*Cas9, altering communication between *Geo*HNH and adjacent RuvC and REC domains.

Anti-CRISPRs have been co-opted as a biological research tool for spatiotemporal control of Cas9 editing through strategic employment of inhibition(29). This work highlights the necessity for atomistic structural and biophysical studies of Acr mechanisms, and that even broad-spectrum inhibitors like AcrIIC1 maintain atomistic subtleties that may differentiate their effect on related Cas systems. We showed that AcrIIC1 attenuates the affinity of *Geo*Cas9 for sgRNA, also demonstrating that timing and order of assembly of the Cas9 RNP are critical parameters for controlling Cas9 editing windows. However, occupancy of Acr in the GeoHNH catalytic site alone is not sufficient to diminish sgRNA affinity, but rather specific contacts between S78 and H582 are required. These findings argue that spatiotemporal regulation of *Geo*HNH by an Acr cannot be explained solely by steric occlusion, rather, specific sidel7chain chemistry must perturb longl7range communication between *Geo*HNH, REC, and RuvC. The stimulation of *ms* motions via saltl7bridge disruption parallel earlier observations of altered *Geo*HNH dynamics, which in prior work extended the thermostability and editing range of *Geo*Cas9, suggesting that natural Acrs and engineered *Geo*Cas9 variants access overlapping allosteric switches, with outcomes dependent on the specific motional landscape.

The potential for a weak secondary AcrIIC1 binding site occupied at high Acr concentrations in NMR titrations and supported by AlphaFold3 modeling is plausible when considering the inhibitor stoichiometries required for *Geo*Cas9 inhibition. Our results suggest that AcrIIC1 inhibition is far less efficient toward preformed *Geo*Cas9 RNPs than toward apo *Geo*Cas9, thus it is possible that evolutionary pressure would select for Acrs capable of binding at multiple sites on a Cas effector. Indeed, *acr* gene expression increases up to 100-fold within the first 20 minutes of infection, effectively flooding the bacterial cell with high concentrations of Acr protein (30). Although exact intracellular Acr concentrations are unknown, overexpression of the *acr* gene and high Acr:Cas9 ratios would favor promiscuous binding of Acrs to many regions of the Cas9 enzyme, including HNH. Such speculation draws some support from the wide diversity of reported Acr mechanisms, including between structurally similar orthologs (22, 29). Further studies are required to unambiguously map additional binding sites for AcrIIC1 on *Geo*Cas9, though it is possible additional weak contacts between the enzyme and inhibitor influence flexibility as a means of accessing the catalytic site.

Together, the integrated structural biophysics presented here argue that even broadl7spectrum Acrs, like AcrIIC1, can tune Cas9 activity in systeml7specific ways, providing a framework for engineering nextl7generation inhibitors that exploit both orthosteric activel7site engagement and targeted manipulation of allosteric networks to achieve precise, contextl7dependent control of genome editing. These data also highlight the critical importance of atomistic studies for continued elucidation of conformational dynamics and allosteric pathways within Cas systems, thereby informing the rational design of controllable Cas9-Acr complexes.

## Materials and Methods

### Protein expression and purification

Separately, the HNH domain of *Geobacillus stearothermophilus* Cas9 (residues 511-662), AcrIIC1 from *N*. *meningitidis* and S78A and C79P AcrIIC1 variants were cloned into pET28a vectors with N-terminal His_6_-tags and TEV protease cleavage sites. The plasmids were transformed into BL21 (DE3) cells (New England BioLabs), expressed, and purified as previously described in LB media(12, 17). Isotopically labeled samples of *Geo*HNH and AcrIIC1 were grown in M9 minimal media containing CaCl_2_, MgSO_4_, and MEM vitamins and supplemented with ^15^N-labeled ammonium chloride and ^13^C-labeled glucose (1.0 g/L and 2.0 g/L, respectively; Cambridge Isotope Laboratories). Cells were grown to an OD_600_ of 0.8–1.0, induced with 1 mM IPTG, and then incubated for a further 4 hr at 37 °C. Cells containing *Geo*HNH were harvested by centrifugation, resuspended in a buffer of 20 mM HEPES, 150 mM KCl, 5 mM imidazole, and 1 mM PMSF at pH 8.0 (lysis buffer), lysed by ultrasonication, and loaded onto a Ni-NTA column equilibrated with lysis buffer. The column was washed with 1 column volume of lysis buffer and *Geo*HNH was eluted with a linear gradient of imidazole from 0.2-250 mM. The His_6_-tag was cleaved by dialyzing the resulting fractions containing *Geo*HNH against a buffer of 20 mM HEPES, 150 mM KCl, and 5 mM imidazole at pH 8.0 overnight at 4 °C with TEV. The sample was further purified on a Hi-Load Superdex 75 size-exclusion column and dialyzed into a buffer of 20 mM HEPES, 80 mM KCl, 1 mM EDTA, 1 mM DTT, 5% glycerol, and 10% (v/v) D_2_O at pH 7.4.

Cells containing AcrIIC1 and variants were also harvested by centrifugation, resuspended in a buffer of 50 mM Tris-HCl, 500 mM NaCl, 1 mM TCEP, 20 mM imidazole, and 1 mM PMSF at pH 7.5 (Acr lysis buffer), lysed by ultrasonication, and loaded onto a Ni-NTA column equilibrated with Acr lysis buffer. The column was washed with 1 column volume of Acr lysis buffer and eluted with a linear gradient of imidazole from 0.2-250 mM. The His_6_-tag was cleaved by dialyzing the resulting fractions containing AcrIIC1 against a buffer of 50 mM HEPES, 150 mM KCl, 5%(v/v) glycerol and 1 mM DTT at pH 7.5 overnight at 4 °C with TEV. The sample was further purified on a Hi-Load Superdex 75 size-exclusion column. AcrIIC1 protein for NMR spectroscopy was exchanged into a buffer of 20 mM HEPES, 80 mM KCl, 1 mM EDTA, 1 mM DTT, and 5% glycerol at pH 7.5 by Amicon centrifugal cell.

The full-length *Geo*Cas9 plasmid was acquired from Addgene (#87700). *Geo*Cas9 was expressed in TB media and was expressed and purified as previously described(13).

### NMR spectroscopy

Two-dimensional ^1^H-^15^N HSQC NMR spectra were collected at 600 MHz and NMR spin relaxation experiments were carried out at 600 and 850 MHz on Bruker Avance NEO and Avance III HD spectrometers, respectively. All experiments were performed at 25 °C. NMR spectra were processed with NMRPipe and analyzed in NMRFAM-SPARKY(31, 32). NMR assignments of *Geo*HNH were obtained from BMRB entry 50887 (14), while those of AcrIIC1 were obtained from BMRB entry 36471 (33).

NMR titrations were performed on a Bruker Avance NEO 850 MHz spectrometer via a series of ^1^H-^15^N TROSY HSQC spectra with increasing ligand (*i.e*. AcrIIC1) concentration. The ^1^H and ^15^N carrier frequencies were set to the water resonance and 120 ppm, respectively. Samples of WT *Geo*HNH were separately titrated with a WT, S78A, or C79P AcrIIC1 until no further spectral perturbations were detected. Combined ^1^H-^15^N NMR chemical shift perturbations were calculated using the method of Bax and coworkers(34).

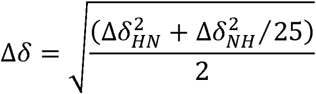

Longitudinal (35) and transverse (36) relaxation rates were measured with randomized *T*_1_ delays of 0, 20, 60, 100, 200, 600, 800, and 1200 ms and *T*_2_ delays of 0, 16.9, 33.9, 50.9, 67.8, 84.8, and 101.8 ms. Peak intensities were quantified in Sparky and the resulting decay profiles were analyzed in Sparky with errors determined from the fitted parameters (32). Uncertainties in these rates were determined from replicate spectra with duplicate relaxation delays of 20 (x2), 60 (x2), 100, 200, 600 (x2), 800, and 1200 ms for *T*_1_ and 16.9, 33.9 (x2), 50.9 (x2), 67.8 (x2), 84.8, 101.8 (x2) ms for *T*_2_. Steady-state ^1^H-[^15^N] NOEs (36) were measured with a 6 s relaxation delay followed by a 3 s saturation (delay) for the saturated (unsaturated) experiments and calculated by I_sat_/I_ref_.

Model-free analysis was conducted via fits to five different forms of the spectral density function using local t_m_, spherical, prolate spheroid, oblate spheroid, or ellipsoid diffusion tensors (37–42). The criteria for inclusion of resonances in the diffusion tensor estimate was based on the method of Bax and coworkers (43). N-H bond lengths were assumed to be 1.02 Å and the ^15^N chemical shift anisotropy tensor was –160 ppm. Diffusion tensor parameters were optimized simultaneously in RELAX under the full automated protocol (44). Model selection was iterated until tensor and order parameters did not deviate from the prior iteration.

CPMG relaxation dispersion experiments were carried out with a constant relaxation period of 20 ms and ν_CPMG_ values of 0, 25, 50 (x2), 75, 100, 150, 250, 500 (x2), 750, 800 (x2), 900, and 1000 Hz (45). Exchange parameters were obtained from fits of the data using the following models:

Model 1: No exchange

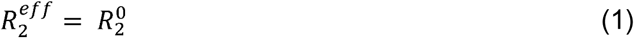

Model 2: Two-state, fast exchange (Meiboom equation (46))

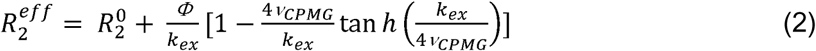

Uncertainties in these rates were determined from replicate spectra with duplicate relaxation delays of 0, 25, 50 (×2), 75, 100, 150, 250, 500 (×2), 750, 800 (×2), 900, and 1000 Hz. All relaxation experiments were carried out in a temperature-compensated interleaved manner.

Histidine ε1 sidechain carbon chemical shifts were measured via HMQC (*hmqcphpr* pulse program) using NMR samples of *Geo*HNH bound to WT or S78A AcrIIC1. The ε1 carbons of the *Geo*HNH histidine sidechains were assigned by mutagenesis to alanine.

### Microscale thermophoresis (MST)

MST experiments were performed on a NanoTemper Monolith X instrument (NanoTemper Technologies) at 25 °C. Experiments quantifying protein-guide RNA interactions used full-length *Geo*Cas9 in complex with WT or S78A AcrIIC1. Signal was detected using a 39 nucleotide Cy5-labeled guide RNA at a concentration of 20 nM in a buffer of 20 mM sodium phosphate, 150 mM KCl, 5 mM MgCl_2_, and 0.1% Triton X-100 at pH 7.6. The *Geo*Cas9 protein was serially diluted from a 115 µM stock into 16 microcentrifuge tubes and combined in a 1:1 molar ratio with serially diluted guide RNA from a 40 nM stock. After incubation for 5 min at 37 °C in the dark, each sample was loaded into an MST capillary (NanoTemper Technologies) for measurement. *K*_d_ values for the various complexes were calculated using the MO Control software (NanoTemper Technologies). Statistical significance was calculated using a two-tailed T-test.

MST experiments quantifying protein-protein interactions used *Geo*Cas9 or *Geo*HNH and WT, S78A, or C79P AcrIIC1. WT AcrIIC1 and its mutants contained 6x-His affinity tags that were used for attachment of Red-tris as a label. The binding affinity of the Red-tris dye for the His-tag was 2 nM. Full-length *Geo*Cas9 or *Geo*HNH were serially diluted from stock concentrations of 95 µM and 200 µM, respectively into microcentrifuge tubes and 100 nM of Red-tris labeled AcrIIC1 was added to each tube. After incubation for 5 min at 37 °C in the dark, each sample was loaded into an MST capillary for measurement. *K*_d_ values for the various complexes were calculated using the MO Control software (NanoTemper Technologies). Statistical significance was calculated using a two-tailed T-test.

### Circular dichroism (CD) spectroscopy

AcrIIC1 proteins were exchanged into a buffer of 20 mM sodium phosphate at pH 7.5 via Amicon centrifugal cell. Samples were then diluted to 1 μM and loaded into a 1 mm quartz cuvette (JASCO Instruments). A CD spectrum was first measured between 200 – 250 nm, after which the sample was progressively heated from 25 – 90 °C in 1.5 °C increments while ellipticity was monitored at 222 and 208 nm. Measurements made on a buffer-only baseline were subtracted from the sample measurements.

### Generation of the *Geo*HNH:AcrIIC1:AcrIIC1 homology model

Models of the protein complexes were generated with the multimer mode of AlphaFold3. Protein sequences for each chain were considered for *Geo*HNH and WT AcrIIC1, with signal peptides and affinity tags removed. All chains were submitted together to specify the intended 2 AcrIIC1:1 *Geo*HNH stoichiometry for the complex. AlphaFold3 was run with its standard pipeline to build multiple sequence alignments and search for structural templates. Both templatel7based and nol7template runs were performed using default settings, unless stated otherwise. For each complex, 5–10 models were generated using different random seeds.

Models were ranked using AlphaFold’s confidence scores, including pLDDT for local perl7residue confidence and combined ipTM/pTM for overall complex and interface quality. The highestl7ranked model was chosen for analysis. Predicted aligned error (PAE) plots and visual inspection in PyMoL were used to assess the reliability of domain and subunit positions; regions with low confidence were interpreted cautiously. Figures of the subsequent AlphaFold structures were generated in PyMoL.

### X-ray Crystallography

*Geo*HNH-AcrIIC1 protein complexes used for crystallization were purified as described above and stored in a crystallization buffer of 20 mM HEPES and 80 mM KCl at pH 8.0. The protein complex was formed by incubation *Geo*HNH and AcrIIC1 proteins at a 1:1 molar ratio for 30 minutes on ice. The complex mixture was then loaded onto an EnrichSEC column and eluted using crystallization buffer. Fractions containing both proteins were used to grow crystals using sitting drop vapor diffusion at room temperature by mixing 0.2 µL of 23 mg/mL *Geo*HNH with 0.4 µL of crystallizing condition from the PEG/Ion (Hampton Research) well E6: containing 0.2 M sodium malonate at pH 6.0 and 20% (w/v) polyethylene glycol 3350. Crystals were looped and frozen using 20% polyethylene glycol as a cryoprotectant. Diffraction images were collected using a Rigaku MicroMax-003i sealed tube X-ray generator with a Saturn 944 HG CCD detector. Images were processed using XDS and Aimless in CCP4 (47, 48). An X-ray structure of AcrIIC1 bound to the *N. meningitidis* Cas9 HNH domain (PDB ID: 8IF0) was used for molecular replacement with Phaser followed by AutoBuild in Phenix (49). The *Geo*HNH-AcrIIC1 structure was finalized through manual building in Coot and refinement in Phenix (49, 50).

### DNA cleavage assays

The inhibition of AcrIIC1 and its variants against *Geo*Cas9 cleavage of DNA substrates was assayed as described previously (14). Briefly, a 1 µL volume of 0, 6, 18, 30, or 60 µM AcrIIC1 was mixed with 6 µM *Geo*Cas9 in a 1x reaction buffer of 20 mM Tris, 100 mM KCl, 5 mM MgCl_2_, 1 mM DTT, and 5% glycerol at pH 7.5 and incubated at 37 °C for 10 min. The mixture was then placed on ice and 1 µL of 6 µM Tnnt2 sgRNA was added along with 1 µL of 140.3 ng/µL PCR-amplified Tnnt2 DNA substrates and 5 µL of 2x reaction buffer. After gentle mixing and a transient spin, the tubes were incubated at 37 °C for 30 min, followed by an incubation at 85 °C for 10 min to terminate the cleavage reaction. Finally, 2 µL of 6x DNA loading buffer was added and 5.5 µL of that reaction mixture was subsequently run on a 1.5% agarose gel. DNA band intensity quantitation was carried out in ImageJ. A negative control reaction without sgRNA and any form of the AcrIIC1 inhibitor was also included.

The sgRNA sequence, with <u>23-nt spacer</u> underlined, was: <u>UUGCACCUACCUUCUGGAUGUAC</u>GUCAUAGUUCCCCUGAGAAAUCAGGGUUACUAUGAU AAGGGCUUUCUGCCUAAGGCAGACUGACCCGCGGCGUUGGGGAUCGCCUGUCGCCCGCUUUUGGCGGGCAUUCCCCAUCCUU

The Tnnt2 DNA sequence, with **PAM** in bold and the 23-nt spacer underlined, was: CAAAGAGCTCCTCGTCCAGTGGGAAGAGAGCTGATCTCATTTGTAAGGAATACCCTCTTCATCCCCCACCCTTGCCATTGATCTATTCATTCCATCTCCATGACAACAGGAAGAGAGGGCCCGGCGTGAGGAGGAGGAGAACAGGAGGAAGGCTGAGGATGAGGCCCGGAAGAAGAAGGCT CTGTCCAACATGATGCAC**TTTGGAGG**GTACATCCAGAAGGTAGGTGCAAAGCAGCATCGGG CACCAGGACACCCCAGTGTATCCTCAAGGCCGCCTTTGCTTGGATCCATGAAGAAATTCCCAACTGCTGTGGCTGAAGTCTAAGGTCTGCTCATGTCTAGCCCCTGAGCTGTCTATCAGCCTGACCATGGCTTCAGTAGGAGGGCTCTGCTGTGTGTGACAGTTAGAACACTAATATGTCTCCAAATTCTGGCTCCCCAAAGGGACAACTGGGAGAATCTTGGGTCCTGGAGTCCAT)

### Molecular dynamics (MD) simulations

All-atom molecular dynamics simulations were performed on the *Geo*HNH-AcrIIC1 complex in its WT form and with three AcrIIC1 variants; S78A, C79P, and S78A/C79P. Initial coordinates were derived from the corresponding X-ray crystal structures of the WT or S78A complexes. Each system consisted of the *Geo*HNH domain bound to AcrIIC1 and was simulated independently in three replicas.

System preparation and simulation protocols were adapted from workflows previously developed and validated for CRISPR–Cas nucleoprotein complexes (23, 51–53). Each complex was solvated in explicit water using rectangular periodic boxes extending at least 12 Å from any solute atom, yielding systems of approximately 32,000 atoms. Appropriate numbers of counterions were added to ensure electrical neutrality and to reproduce physiological salt conditions. All simulations were carried out using the AMBER ff19SB19 force field for proteins, together with OL15 and OL3 corrections for DNA and RNA, respectively. Solvent molecules were represented using the TIP3P water model (54–57). Hydrogen-containing bonds were constrained using the LINCS algorithm, enabling a 2 fs integration time step. Long-range electrostatic interactions were evaluated using the particle mesh Ewald method with a real-space cutoff of 10 Å, and van der Waals interactions were truncated at the same distance. Energy minimization was first performed to eliminate unfavorable contacts while restraining the protein backbone with harmonic potentials of 100 kcal·mol⁻¹·Å⁻². Systems were then gradually heated from 0 K to 300 K under constant-volume conditions, followed by equilibration in the isothermal–isobaric ensemble. Temperature regulation was achieved using Langevin dynamics with a collision frequency of 1 ps⁻¹, and pressure was maintained at 1 atm using the Berendsen barostat (58, 59). After equilibration, restraints were removed and production simulations were carried out in the NVT ensemble. Each system was simulated for 2 μs per replica, with three independent replicas, yielding 6 μs of sampling per variant and a total simulation time of 24 μs. Equations of motion were propagated using the leapfrog Verlet algorithm. All simulations were performed using the GPU-accelerated version of AMBER 22 (60). Trajectory analyses were conducted on the combined ensembles using standard AMBER tools and custom scripts.

To characterize differences in the stability of *Geo*HNH-AcrIIC1 intermolecular contacts across systems, residue-residue contact maps were first computed for each trajectory. A contact between two residues was defined as formed when any pair of heavy atoms were within a distance cutoff of ≤4.5 Å. For each residue pair, a contact persistence frequency (*f*) was calculated as the fraction of trajectory frames in which the contact was present relative to the total number of analyzed frames. To enable a direct comparison between systems, only *common contacts* (those observed at least once in both the WT and variants) were considered. The difference in contact stability was then quantified as:

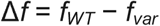

where *f_WT_* and *f_var_* represent the contact persistence frequencies in the WT *Geo*HNH-AcrIIC1 and mutant *Geo*HNH-AcrIIC1 complexes, respectively. Positive values of Δ*f* indicate contacts that are more stable in the WT system, whereas negative values indicate contacts that are more stable in the variant.

### Binding free energy calculations

The binding free energy (Δ*G*_bind_) between *Geo*HNH and AcrIIC1 was calculated using the Molecular Mechanics Generalized Born Surface Area (MM-GBSA) method. For each system, the average binding energy (Δ*G*_bind_) values were calculated from the independent windows chosen at a regular interval of 2 ns during the 20 ns of the stable trajectory. The binding free energy was obtained from the difference between the free energies of the protein-protein complex (*G*_complex_), unbound protein (*G*_HNH_) and the protein (*G*_AcrIIC1_):

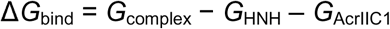

In Amber, these binding energy values are computed from the changes in the molecular mechanical energy (Δ*E*_MM_), entropic contribution, and the change in solvation free energy due to the substrate binding to the protein for complex formation:

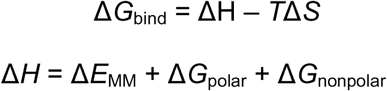

The term (Δ*E*_MM_) includes electrostatic and van der Waals interactions between the substrate and protein. The Δ*G*_polar_ is estimated by using the Generalized born (GB) model and the Δ*G*_nonpolar_ is obtained from the hydrophobic contributions calculated from the solvent accessible surface area. To identify sequence-specific contributions, MM/GBSA energy decomposition was performed at the per-base level.

## Acknowledgments

This work was supported by NIH grant R01 GM136815 (to GP and GPL) and NSF grant (MCB 2143760 to GPL). ALK was supported by a Blavatnik Family Graduate Fellowship in Biology & Medicine.

## Data Availability

X-ray crystal structures of *Geo*HNH bound to WT AcrIIC1 and S78A AcrIIC1 were deposited under PDB IDs 10VC and 10VB, respectively. Data or resources related to this work are available from the corresponding authors upon request.

## References

1. Jinek M, Chylinski K, Fonfara I, Hauer M, Doudna JA, Charpentier E. A Programmable Dual-RNA–Guided DNA Endonuclease in Adaptive Bacterial Immunity. Science. 2012;337(6096):816–21.

2. Chen K, Han H, Zhao S, Xu B, Yin B, Lawanprasert A, et al. Lung and liver editing by lipid nanoparticle delivery of a stable CRISPR–Cas9 ribonucleoprotein. Nature Biotechnology. 2025;43(9):1445–57.

3. Li T, Yang Y, Qi H, Cui W, Zhang L, Fu X, et al. CRISPR/Cas9 therapeutics: progress and prospects. Signal Transduction and Targeted Therapy. 2023;8(1).

4. Zhang S, Wang Y, Mao D, Wang Y, Zhang H, Pan Y, et al. Current trends of clinical trials involving CRISPR/Cas systems. Frontiers in Medicine. 2023;10.

5. Sousa AA, Terrey M, Sakai HA, Simmons CQ, Arystarkhova E, Morsci NS, et al. In vivo prime editing rescues alternating hemiplegia of childhood in mice. Cell. 2025;188(16):4275–94.e23.

6. Anastassopoulou C, Tsakri D, Panagiotopoulos A-P, Saldari C, Sagona AP, Tsakris A. Armed Phages: A New Weapon in the Battle Against Antimicrobial Resistance. Viruses. 2025;17(7):911.

7. Fu Y, Foden JA, Khayter C, Maeder ML, Reyon D, Joung JK, et al. High-frequency off-target mutagenesis induced by CRISPR-Cas nucleases in human cells. Nature Biotechnology. 2013;31(9):822–6.

8. Banani SF, Lee HO, Hyman AA, Rosen MK. Biomolecular condensates: organizers of cellular biochemistry. Nat Rev Mol Cell Biol. 2017;18(5):285–98.

9. Chen JS, Dagdas YS, Kleinstiver BP, Welch MM, Sousa AA, Harrington LB, et al. Enhanced proofreading governs CRISPR–Cas9 targeting accuracy. Nature. 2017;550(7676):407–10.

10. Casini A, Olivieri M, Petris G, Montagna C, Reginato G, Maule G, et al. A highly specific SpCas9 variant is identified by in vivo screening in yeast. Nature Biotechnology. 2018;36(3):265–71.

11. Kleinstiver BP, Pattanayak V, Prew MS, Tsai SQ, Nguyen NT, Zheng Z, et al. High-fidelity CRISPR–Cas9 nucleases with no detectable genome-wide off-target effects. Nature. 2016;529(7587):490–5.

12. Belato HB, Norbrun C, Luo J, Pindi C, Sinha S, D’Ordine AM, et al. Disruption of electrostatic contacts in the HNH nuclease from a thermophilic Cas9 rewires allosteric motions and enhances high-temperature DNA cleavage. The Journal of Chemical Physics. 2022;157(22):225103.

13. Harrington LB, Paez-Espino D, Staahl BT, Chen JS, Ma E, Kyrpides NC, et al. A thermostable Cas9 with increased lifetime in human plasma. Nature Communications. 2017;8(1).

14. Belato HB, Knight AL, D’Ordine AM, Pindi C, Fan Z, Luo J, et al. Structural and dynamic impacts of single-atom disruptions to guide RNA interactions within the recognition lobe of Geobacillus stearothermophilus Cas9. eLife. 2025;13.

15. Eggers AR, Chen K, Soczek KM, Tuck OT, Doherty EE, Xu B, et al. Rapid DNA unwinding accelerates genome editing by engineered CRISPR-Cas9. Cell. 2024;187(13):3249–61.e14.

16. Shen P, Zhang L, Liu B, Li X, Wang C, Min J, et al. Structure of *Geobacillus stearothermophilus* Cas9: Insights into the Catalytic Process and Thermostability of CRISPR-Cas9. ACS Catalysis. 2024;14(17):13227–35.

17. Harrington LB, Doxzen KW, Ma E, Liu J-J, Knott GJ, Edraki A, et al. A Broad-Spectrum Inhibitor of CRISPR-Cas9. Cell. 2017;170(6):1224–33.e15.

18. Zheng I, Learn B, Bailey S. Structural basis for inhibition of SpyCas9 by the anti-CRISPR protein AcrIIA26. Biochemical Journal. 2026;483(3):289–300.

19. Kang YW, Park HH. The anti– <scp>CRISPR</scp> protein <scp>AcrIE8</scp>.1 inhibits the type I-E <scp>CRISPR</scp> –Cas system by directly binding to the. FEBS Letters. 2025.

20. Yu L, Yin M, Zhu Y, Lu Z, Xiao B, Zhou F, et al. An anti-CRISPR targets the sgRNA to block Cas9 and guides the design of enhanced genome editors. Nature Structural & Molecular Biology. 2026.

21. Kim Y, Lee SJ, Yoon HJ, Kim NK, Lee BJ, Suh JY. Anti-CRISPR AcrIIC3 discriminates between Cas9 orthologs via targeting the variable surface of the HNH nuclease domain. The FEBS Journal. 2019;286(23):4661–74.

22. Zhu Y, Gao A, Zhan Q, Wang Y, Feng H, Liu S, et al. Diverse Mechanisms of CRISPR-Cas9 Inhibition by Type IIC Anti-CRISPR Proteins. Molecular Cell. 2019;74(2):296–309.e7.

23. Skeens E, Sinha S, Ahsan M, D’Ordine AM, Jogl G, Palermo G, et al. High-fidelity, hyper-accurate, and evolved mutants rewire atomic-level communication in CRISPR-Cas9. Science Advances. 2024;10(10).

24. Wang J, Maschietto F, Qiu T, Arantes PR, Skeens E, Palermo G, et al. Substrate-independent activation pathways of the CRISPR-Cas9 HNH nuclease. Biophysical Journal. 2023;122(24):4635–44.

25. Wang J, Arantes PR, Ahsan M, Sinha S, Kyro GW, Maschietto F, et al. Twisting and swiveling domain motions in Cas9 to recognize target DNA duplexes, make double-strand breaks, and release cleaved duplexes. Frontiers in Molecular Biosciences. 2023;9.

26. Belato HB, D’Ordine AM, Nierzwicki L, Arantes PR, Jogl G, Palermo G, et al. Structural and dynamic insights into the HNH nuclease of divergent Cas9 species. Journal of Structural Biology. 2022;214(1):107814.

27. Abramson J, Adler J, Dunger J, Evans R, Green T, Pritzel A, et al. Accurate structure prediction of biomolecular interactions with AlphaFold 3. Nature. 2024;630(8016):493–500.

28. Zhu YL. NmeHNH-AcrIIC1 Complex Protein Data Bank, PDB. 2024.

29. Marino ND, Pinilla-Redondo R, Csörgő B, Bondy-Denomy J. Anti-CRISPR protein applications: natural brakes for CRISPR-Cas technologies. Nature Methods. 2020;17(5):471–9.

30. Li Y, Bondy-Denomy J. Anti-CRISPRs go viral: The infection biology of CRISPR-Cas inhibitors. Cell Host & Microbe. 2021;29(5):704–14.

31. Delaglio F, Grzesiek S, Vuister G, Zhu G, Pfeifer J, Bax A. NMRPipe: A multidimensional spectral processing system based on UNIX pipes. Journal of Biomolecular NMR. 1995;6(3).

32. Lee W, Tonelli M, Markley JL. NMRFAM-SPARKY: enhanced software for biomolecular NMR spectroscopy. Bioinformatics. 2015;31(8):1325–7.

33. Zhao Y, Hu J, Yang S-S, Zhong J, Liu J, Wang S, et al. A redox switch regulates the assembly and anti-CRISPR activity of AcrIIC1. Nature Communications. 2022;13(1).

34. Grzesiek S, Stahl SJ, Wingfield PT, Bax A. The CD4 Determinant for Downregulation by HIV-1 Nef Directly Binds to Nef. Mapping of the Nef Binding Surface by NMR. Biochemistry. 1996;35(32):10256–61.

35. Lakomek N-A, Ying J, Bax A. Measurement of 15N relaxation rates in perdeuterated proteins by TROSY-based methods. Journal of Biomolecular NMR. 2012;53(3):209–21.

36. Zhu G, Xia Y, Nicholson LK, Sze KH. Protein Dynamics Measurements by TROSY-Based NMR Experiments. Journal of Magnetic Resonance. 2000;143(2):423–6.

37. Brüschweiler R, Liao X, Wright PE. Long-range Motional Restrictions in a Multidomain Zinc-finger Protein from Anisotropic Tumbling. Science. 1995;268:886–9.

38. Mandel AM, Akke M, Palmer AG. Backbone Dynamics of Escherichia coli Ribonuclease HI: Correlations with Structure and Function in an Active Enzyme. J Mol Biol. 1995;246:144–63.

39. Fushman D, Cahill S, Cowburn D. The Main-chain Dynamics of the Dynamin Pleckstrin Homology (PH) Domain in Solution: Analysis of 15N Relaxation with Monomer/Dimer Equilibration. J Mol Biol. 1997;266:173–94.

40. Orekhov VY, Korzhnev DM, Diercks T, Kessler H, Arseniev AS. 1H-15N NMR Dynamic Study of an Isolated α-helical Peptide (1-36)bacteriorhodopsin Reveals the Equilibrium Helix-coil Transitions. J Biomol NMR. 1999;14:345–56.

41. Korzhnev DM, Billeter M, Arseniev AS, Orekhov VY. NMR Studies of Brownian Tumbling and Internal Motions in Proteins. Prog NMR Spectros. 2001;38:197–266.

42. Zhuravleva AV, Korzhnev DM, Kupce E, Arseniev AS, Billeter M, Orekhov VY. Gated Electron Transfers and Electron Pathways in Azurin: A NMR Dynamic Study at Multiple Fields and Temperatures. J Mol Biol. 2004;342:1599–611.

43. Tjandra N, Feller SE, Pastor RW, Bax A. Rotational Diffusion Anisotropy of Human Ubiquitin from N-15 NMR Relaxation. J Am Chem Soc. 1995;117:12562–6.

44. Bieri M, d’Auvergne EJ, Gooley PR. relaxGUI: a new software for fast and simple NMR relaxation data analysis and calculation of ps-ns and μs motion of proteins. J Biomol NMR. 2011;50(2):147–55.

45. Loria JP, Rance M, Palmer AG. A Relaxation-Compensated Carr−Purcell−Meiboom−Gill Sequence for Characterizing Chemical Exchange by NMR Spectroscopy. Journal of the American Chemical Society. 1999;121(10):2331–2.

46. Luz Z, Meiboom S. Nuclear Magnetic Resonance Study of the Protolysis of Trimethylammonium Ion in Aqueous Solution—Order of the Reaction with Respect to Solvent. The Journal of Chemical Physics. 1963;39(2):366–70.

47. Kabsch W. XDS. Acta Crystallographica Section D Biological Crystallography. 2010;66(2):125–32.

48. Winn MD, Ballard CC, Cowtan KD, Dodson EJ, Emsley P, Evans PR, et al. Overview of the*CCP*4 suite and current developments. Acta Crystallographica Section D Biological Crystallography. 2011;67(4):235–42.

49. Liebschner D, Afonine PV, Baker ML, Bunkóczi G, Chen VB, Croll TI, et al. Macromolecular structure determination using X-rays, neutrons and electrons: recent developments in *Phenix*. Acta Crystallographica Section D Structural Biology. 2019;75(10):861–77.

50. Emsley P, Lohkamp B, Scott WG, Cowtan K. Features and development of *Coot*. Acta Crystallographica Section D Biological Crystallography. 2010;66(4):486–501.

51. Patel AC, Sinha S, Arantes PR, Palermo G. Unveiling Cas8 dynamics and regulation within a transposon-encoded Cascade–TniQ complex. Proceedings of the National Academy of Sciences. 2025;122(14).

52. Sinha S, Adrian, Pablo, Patel A, Mitchell, Palermo G. Unveiling the RNA-mediated allosteric activation discloses functional hotspots in CRISPR-Cas13a. Nucleic Acids Research. 2024;52(2):906–20.

53. Pablo, Chen X, Sinha S, Saha A, Amun, Sample M, et al. Dimerization of the deaminase domain and locking interactions with Cas9 boost base editing efficiency in ABE8e. Nucleic Acids Research. 2024;52(22):13931–44.

54. Tian C, Kasavajhala K, Belfon KAA, Raguette L, Huang H, Migues AN, et al. ff19SB: Amino-Acid-Specific Protein Backbone Parameters Trained against Quantum Mechanics Energy Surfaces in Solution. Journal of Chemical Theory and Computation. 2020;16(1):528–52.

55. Galindo-Murillo R, Robertson JC, Zgarbová M, Šponer J, Otyepka M, Jurečka P, et al. Assessing the Current State of Amber Force Field Modifications for DNA. Journal of Chemical Theory and Computation. 2016;12(8):4114–27.

56. Zgarbová M, Otyepka M, Šponer J, Mládek A, Banáš P, Cheatham TE, et al. Refinement of the Cornell et al. Nucleic Acids Force Field Based on Reference Quantum Chemical Calculations of Glycosidic Torsion Profiles. Journal of Chemical Theory and Computation. 2011;7(9):2886–902.

57. Jorgensen WL, Chandrasekhar J, Madura JD, Impey RW, Klein ML. Comparison of simple potential functions for simulating liquid water. The Journal of Chemical Physics. 1983;79(2):926–35.

58. Turq P, Lantelme F, Friedman HL. Brownian dynamics: Its application to ionic solutions. The Journal of Chemical Physics. 1977;66(7):3039–44.

59. Berendsen HJC, Postma JPM, Van Gunsteren WF, Dinola A, Haak JR. Molecular dynamics with coupling to an external bath. The Journal of Chemical Physics. 1984;81(8):3684–90.

60. Case DA, Aktulga HM, Belfon K, Cerutti DS, Cisneros GA, Cruzeiro VWD, et al. AmberTools. Journal of Chemical Information and Modeling. 2023;63(20):6183–91.

