## Supplementary Information for "Orthosteric and allosteric effects of anti-CRISPR II-C1 inhibition on *Geo*Cas9 from integrated structural biophysics"



**Figure S1.** Comparison of *NmeCas9* and *GeoCas9* structures bound to AcrIIC1

**Figure S2.** NMR titration of $^{15}$N labeled AcrIIC1 with *Geo*HNH

**Figure S3.** $^{1}$H-$^{15}$N HSQC spectra and CD profiles of AcrIIC1 variants

**Figure S4.** $^{1}$H-$^{15}$N HSQC spectra of *Geo*HNH bound to WT, S78A, and C79P AcrIIC1

**Figure S5.** $T_1$, $T_2$, $^{1}$H-[$^{15}$N]-NOE relaxation parameters of *Geo*HNH in the absence and presence of WT S78A, and C79P AcrIIC1

**Figure S6.** $T_1$, $T_2$, $^{1}$H-[$^{15}$N]-NOE relaxation parameters of *Geo*HNH in the absence and presence of WT AcrIIC1, collected at 600 and 850 MHz for Model-free analysis

**Figure S7.** DNA cleavage assay informing the order of AcrIIC1 and sgRNA binding in *Geo*Cas9 function

**Figure S8.** MD derived mutation-induced changes *Geo*HNH-AcrIIC1 complex.

**Figure S9.** Structure of sgRNA:DNA bound *Geo*Cas9 compared to that of *Geo*HNH:AcrIIC1, examining steric clash of domains

**Figure S10.** Representative MST binding curves testing the effect of WT and S78A AcrIIC1 on the affinity of *Geo*Cas9 for sgRNA

**Figure S11.** Representative MST binding curves measuring the affinity of *Geo*HNH for WT and S78A AcrIIC1

**Table S1.** Crystallographic data reported for structures of WT and S78A AcrIIC1 bound to *Geo*HNH

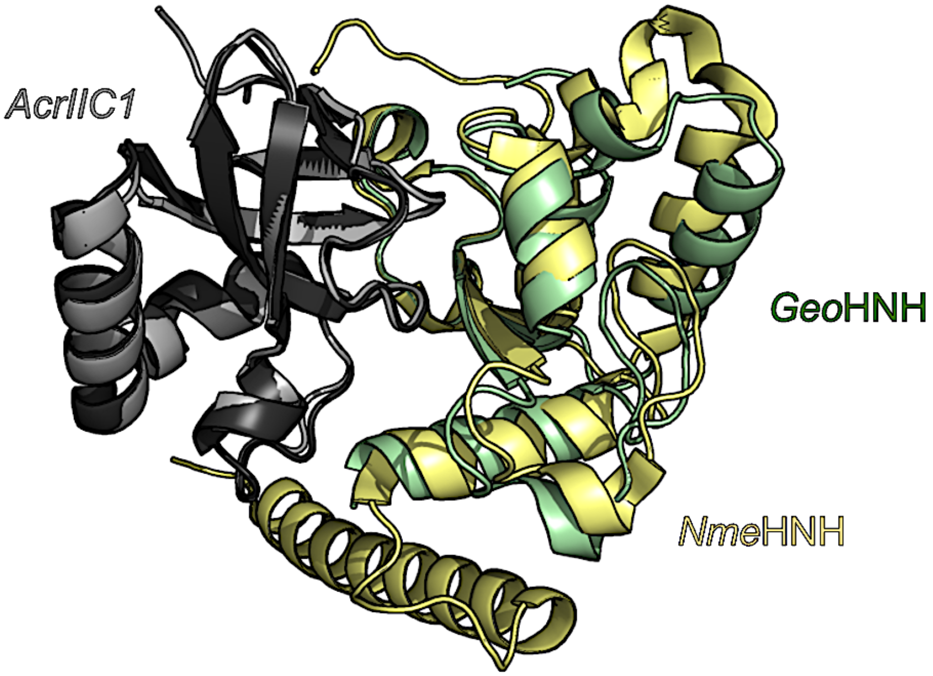


**Figure S1.** Overlay of X-ray crystal structures of AcrIIC1 bound to *Nme*HNH (PDB ID: 5VGB) and to *Geo*HNH (PDB ID: 10VC). A very high degree of similarity is noted.

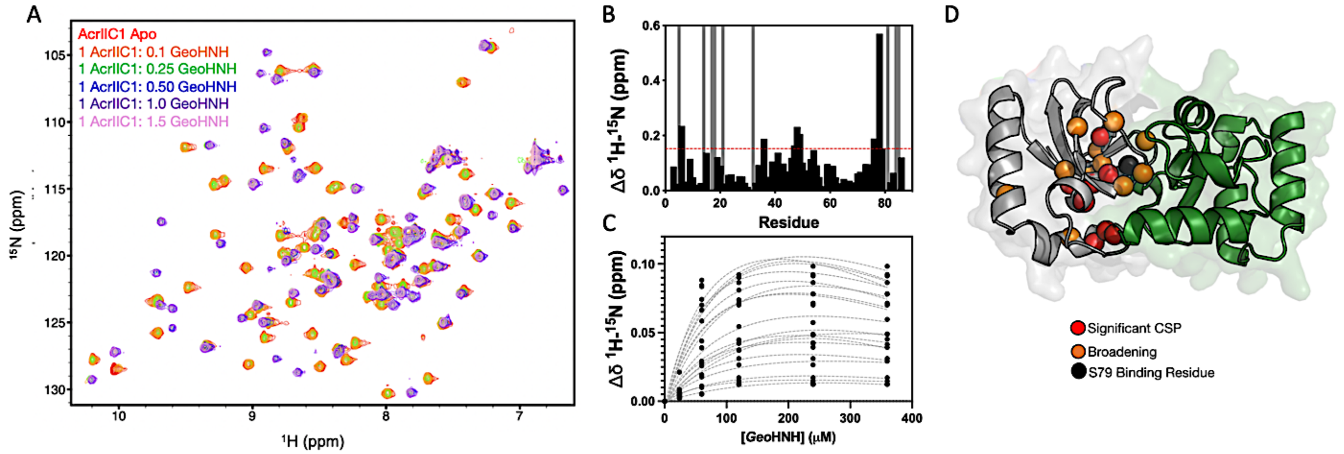


**Figure S2. (A)** $^{1}$H-$^{15}$N HSQC spectral titration of $^{15}$N-AcrIIC1 with *Geo*HNH. **(B)** Per-residue CSPs within AcrIIC1 caused by *Geo*HNH. Grey bars denote sites of NMR line broadening. Red dashed vertical line are CSPs >1.5$\sigma$ of the 10% trimmed mean of all shifts. **(C)** Apparent binding affinity of AcrIIC1 for *Geo*HNH, based on a global fit of fast-exchange NMR chemical shift trajectories, is ~15 $\mu$M. **(D)** CSPs >1.5$\sigma$ of the 10% trimmed mean of all shifts (horizontal dashed line in (B)) are mapped onto the AcrIIC1 structure as spheres, where red denotes CSPs >1.5$\sigma$, orange denotes line broadening and black denotes the binding site residue S78.

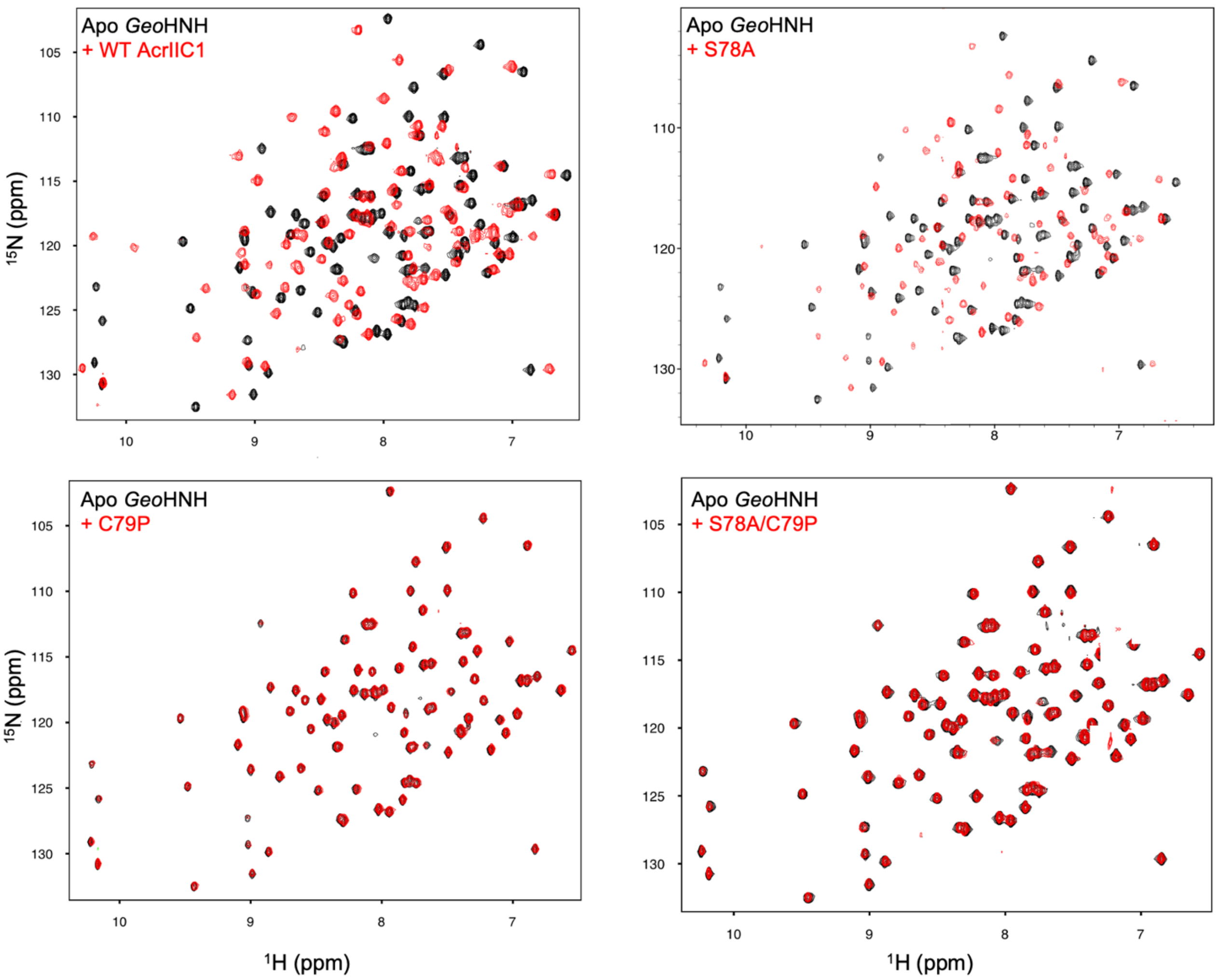


**Figure S3.** $^{1}$H-$^{15}$N HSQC NMR spectra of *Geo*HNH titrated with WT, S78A, C79P, or S78A/C79P AcrIIC1. C79P and S78A/C79P AcrIIC1 inhibitors do not induce any CSPs in *Geo*HNH.

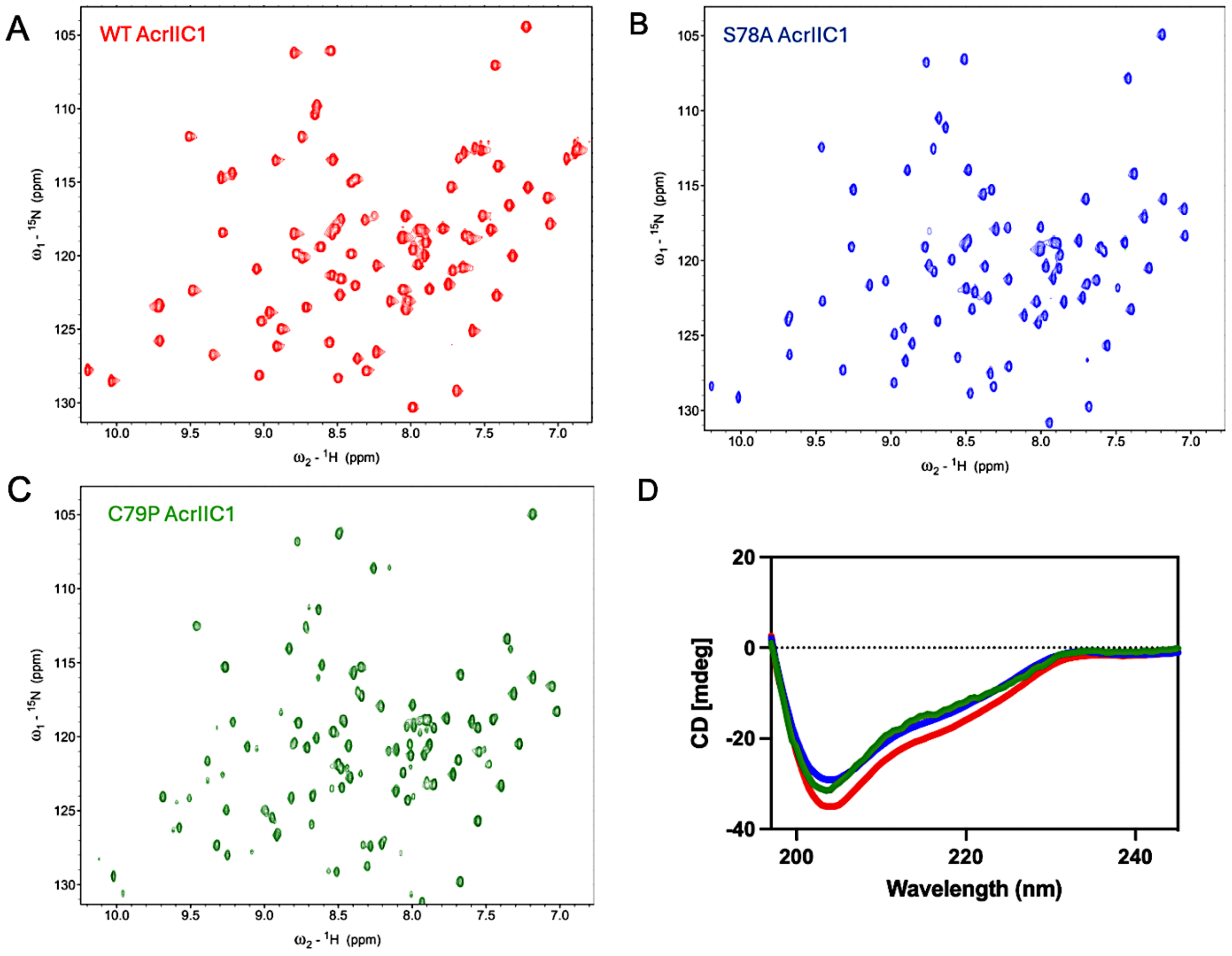


**Figure S4.** $^{1}H$-$^{15}N$ HSQC spectra of **(A)** WT AcrIIC1 **(B)** S78A AcrIIC1 and **(C)** C79P AcrIIC1. **(D)** CD spectra of WT (red), S78A (blue) C79P AcrIIC1 (green) showing maintenance of secondary structure in the presence of mutations.

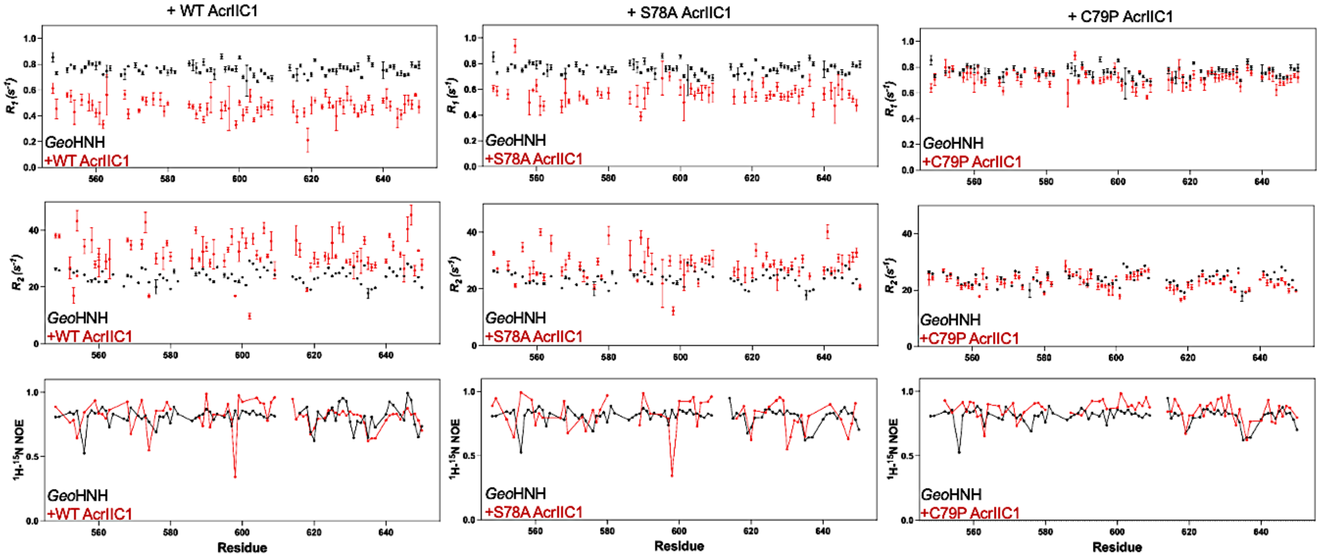


**Figure S5.** Summary of $R_1$, $R_2$, and $^1$H-[$^{15}$N] heteronuclear NOE relaxation parameters. The per-residue $R_1$, $R_2$, and NOE values are plotted for *Geo*HNH in the presence of WT (left), S78A (center), and C79P (right) AcrIIC1.

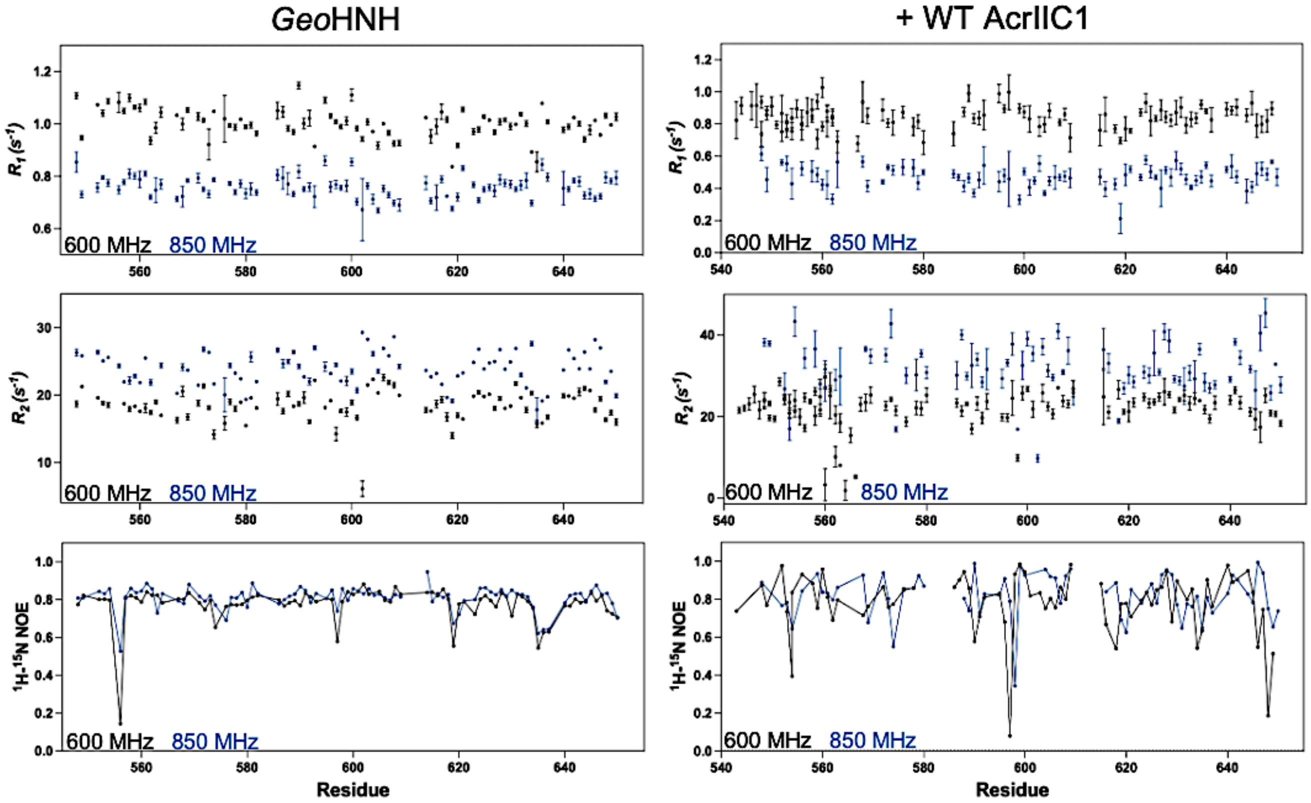


**Figure S6.** Summary of $R_1$, $R_2$, and $^1$H-[$^{15}$N] heteronuclear NOE relaxation parameters collected at 600 and 850 MHz. The per-residue $R_1$, $R_2$, and NOE values are plotted for apo and WT AcrIIC1-bound *Geo*HNH.

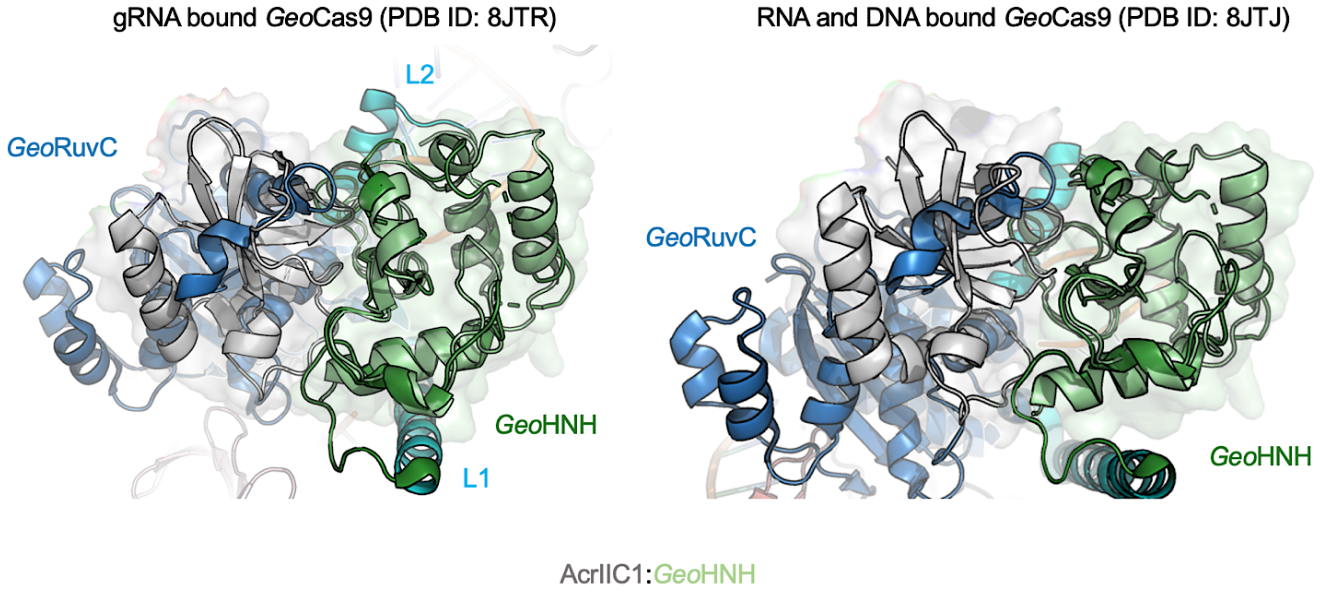


**Figure S7.** Structures of sgRNA-bound (left) and sgRNA:DNA-bound *Geo*Cas9 revealing potential steric clash of AcrIIC1 with RuvC when these structures are overlaid with the *Geo*HNH-AcrIIC1 crystal structure.

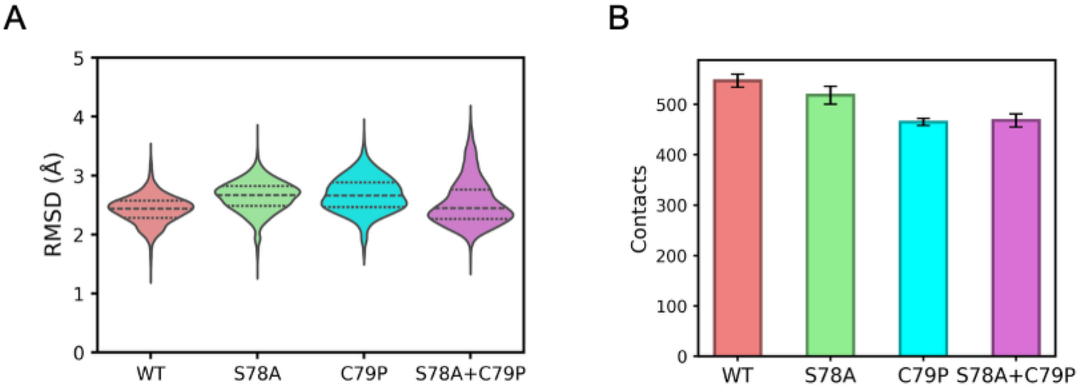


**Figure S8.** Mutation-dependent changes GeoHNH-AcrIIC1 complex. A) Backbone RMSD distributions from triplicate MD simulations for WT and mutant complexes (S78A, C79P, and S78A/C79P). B) Average number of intermolecular contacts between GeoHNH and AcrIIC1 in WT and mutant systems, defined using a 4.5 Å heavy-atom distance cutoff. Error bars represent the standard error of the mean (SEM) calculated across three independent 2 μs trajectories for each system.

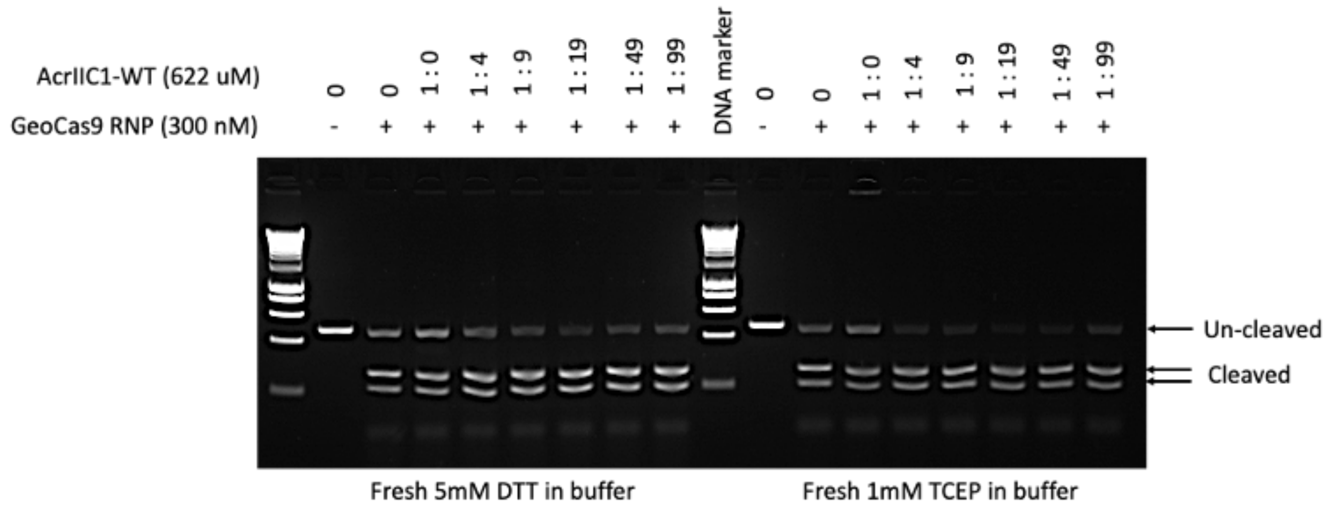


**Figure S9.** Representative gel monitoring *in vitro* DNA cleavage. If the *Geo*Cas9 RNP is formed _prior to the addition of AcrIIC1_, a significant reduction in the inhibiton efficacy of AcrIIC1 is observed. Thus, while AcrIIC1 can inhibit *Geo*Cas9 at higher concentrations, it is most effective if it binds *Geo*Cas9 prior to sgRNA.

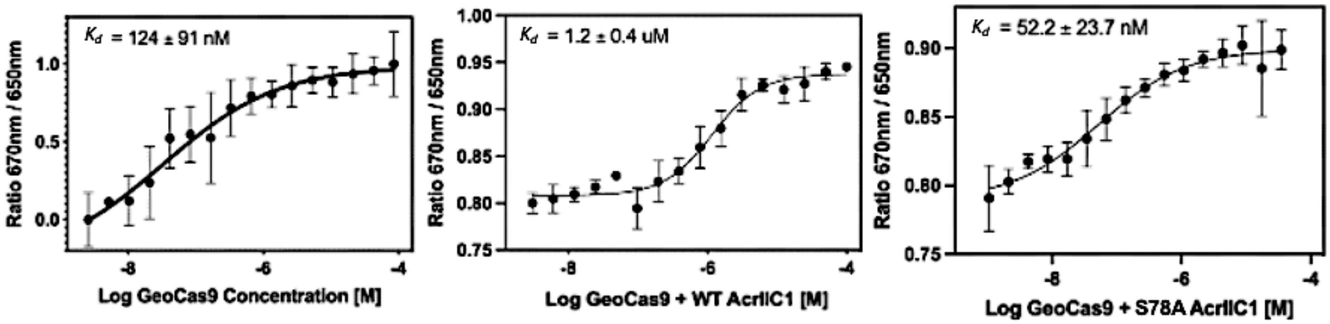


**Figure S10.** Representative MST profiles monitoring *Geo*Cas9, *Geo*Cas9 + WT AcrIIC1, and *Geo*Cas9 + S78A AcrIIC1 binding to a Cy5-labeled 8UZA gRNA. $K_d$ values across n≥3 technical replicates are inset in each plot.

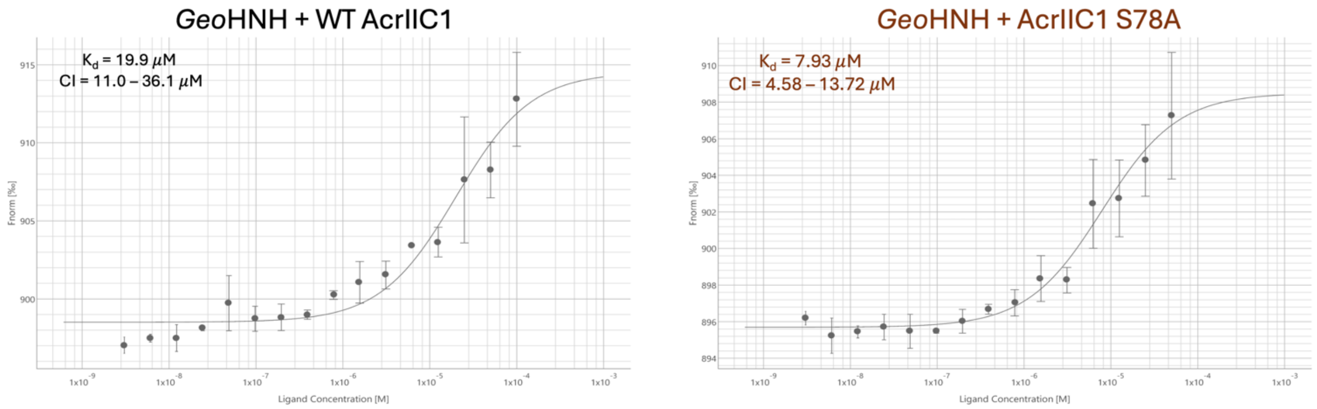


**Figure S11.** Representative MST profiles monitoring *Geo*HNH binding to a fluorescently labeled WT AcrIIC1 or S78A AcrIIC1. $K_d$ values across n≥3 technical replicates are inset in each plot.

**Table S1.** Parameters of crystallographic data collection, refinement, and validation.

| PDB ID | **10VC** | **10VB** |
|---|---|---|
| **Mutation(s)** | | S78A |
| **Data collection**[1] | | |
| Space group | P1 $2_1$ 1 | P1 $2_1$ 1 |
| Cell dimensions | | |
| $\alpha$, $\beta$, $\gamma$ (°) | 90, 93.65, 90 | 90, 93.82, 90 |
| Resolution (Å)[2] | 29.59 – 1.79 (1.81 – 1.79) | 45.23 – 2.30 (2.34 – 2.30) |
| $CC_{1/2}$ | 0.997 (0.42) | 0.991 (0.38) |
| $I / \sigma I$ | 2.50 (2.03) | 3.10 (2.55) |
| Completeness (%) | 99.95 (100.00) | 97.99 (75.21) |
| Multiplicity | 5.7 | 3.7 |
| **Refinement** | | |
| Resolution (Å) | 29.59 – 1.79 (1.81 – 1.79) | 45.23 – 2.301 (2.34 – 2.3) |
| Reflections | 119809 (3995) | 56470 (2069) |
| $R_{work}$ / $R_{free}$ | 0.1985/0.2273 (0.2620/0.2997) | 0.2092/0.2791 (0.2393/0.3433) |
| **Model composition** | | |
| Non-hydrogen atoms | 10744 | 10194 |
| macromolecules | 9450 | 9444 |
| Average B factor | 23.61 | 24.65 |
| Protein residues | 1159 | 1159 |
| **Root mean square deviations** | | |
| Bond lengths (Å) | 0.015 | 0.009 |
| Bond angles (°) | 1.43 | 0.96 |
| **Validation** | | |
| Clashscore | 10.93 | 9.21 |
| Ramachandran plot (%) | | |
| Favored | 97.62 | 96.48 |
| Allowed | 2.2 | 3.17 |
| Outliers | 0.18 | 0.35 |
| Rotamer outliers (%) | 0.5 | 1.91 |

[1]One crystal was used for each data set.

[2]Highest resolution shell is shown in parenthesis.
